# Modulation of aggression by social novelty recognition memory in the hippocampal CA2 region

**DOI:** 10.1101/2024.05.03.592403

**Authors:** Andres Villegas, Steven A. Siegelbaum

## Abstract

The dorsal CA2 subregion (dCA2) of the hippocampus exerts a critical role in social novelty recognition (SNR) memory and in the promotion of social aggression. Whether the social aggression and SNR memory functions of dCA2 are related or represent independent processes is unknown. Here we investigated the hypotheses that an animal is more likely to attack a novel compared to familiar animal and that dCA2 promotes social aggression through its ability to discriminate between novel and familiar conspecifics. To test these ideas, we conducted a multi-day resident intruder (R-I) test of aggression towards novel and familiar conspecifics. We found that mice were more likely to attack a novel compared to familiarized intruder and that silencing of dCA2 caused a more profound inhibition of aggression towards a novel than familiarized intruder. To explore whether and how dCA2 pyramidal neurons encode aggression, we recorded their activity using microendoscopic calcium imaging throughout the days of the R-I test. We found that a fraction of dCA2 neurons were selectively activated or inhibited during exploration, dominance, and attack behaviors and that these signals were enhanced during interaction with a novel compared to familiarized conspecific. Based on dCA2 population activity, a set of binary linear classifiers accurately decoded whether an animal was engaged in each of these forms of social behavior. Of particular interest, the accuracy of decoding aggression was greater with novel compared to familiarized intruders, with significant cross-day decoding using the same familiar animal on each day but not for a familiar-novel pair. Together, these findings demonstrate that dCA2 integrates information about social novelty with signals of behavioral state to promote aggression towards novel conspecifics.

## Introduction

Social memory plays a crucial role in a range of adaptive social behaviors, enabling individuals to build and refine responses that are contextually appropriate based on prior social experience. The formation of a social memory begins with the discrimination of a novel from familiar conspecific and is further modified by ongoing social experience. Adult mice have an innate preference for social exploration of a novel relative to a familiar mouse^1^, which serves as a proxy for social novelty recognition (SNR) memory, and, accordingly, attenuates after sequential encounters with the same conspecific^2–4^. However, whether and how social memory influences specific innate behaviors remains unknown.

One salient innate behavior is social aggression. Adaptive aggression is crucial for competitive survival, mating, and formation of stable and healthy social groups. During an encounter between two male conspecifics, an instinctual response is triggered to confront a perceived threat^5^. After initial assessment during mutual social exploration, a sequence of escalating aggressive behavioral states can lead to one animal engaging first in dominance behaviors followed by aggressive attack^6^. The decision to escalate to an attack is a consequence of internal states and external sensory cues, including context and motivation of the two conspecifics^6–8^. Previous studies have demonstrated increased levels of aggression in an animal following repeated fighting experiences, modified by both neuromodulation through social peptides vasopressin and oxytocin and by changes in plasticity at canonical aggression circuits^9–11^. One potential external trigger of aggression is social novelty. Instability in social groups following the introduction of a novel conspecific leads to periods of aggressive outbreaks^12–16^. Whether and how social memory and the detection of social novelty contributes to adaptive aggression, to our knowledge, remains unknown.

The dorsal CA2 subregion (dCA2) of the hippocampus is crucial for the encoding, consolidation, and recall of social memory through a core dCA2 to ventral CA1 (vCA1) circuit^2–4,17^. Recent studies have demonstrated that dCA2 also serves to promote social aggression. Thus, arginine vasopressinergic inputs to dCA2 from the paraventricular nucleus (PVN) of the hypothalamus enhance aggression by acting on the dCA2-enriched arginine vasopressin 1b (Avpr1b) receptor^18–21^. dCA2 activation promotes aggression through its projections to the lateral septum (LS)^22^, which disinhibit the ventral lateral portion of the ventral medial hypothalamus (VMHvl), a region known to promote aggression^23^, providing a top-down control of aggression through a social memory center. Recent findings have provided insight into how dCA2 neural activity encodes conspecific identity and social novelty^24–26^. Furthermore, dCA2 has been found to incorporate both positive^27^ and negative^28^ valence associations into representations of social identity. However, an understanding of the specific neural mechanisms linking SNR memory and social aggression in dCA2 as either independent or complementary processes is lacking. Here we test the hypothesis that dCA2 promotes aggression by providing a social novelty signal, enhancing aggression towards novel compared to familiar conspecifics, by comparing resident aggression towards novel and familiarized intruders in a resident-intruder test (R-I)^29^. We used pharmacogenetic silencing and calcium imaging of dCA2 pyramidal neurons in freely moving resident mice to assess the causal role of dCA2 pyramidal neurons in the display of aggression and related social behaviors towards novel and familiarized intruders.

We find that dCA2 integrates social information of conspecific novelty to guide an adaptive behavioral response to future conflict. Specifically, dCA2 promotes aggression dependent on the novelty of a conspecific and prior social experience. This behavioral role of dCA2 is reflected in differential population-level encoding of aggression-related behaviors towards novel and familiar conspecifics. Our study thereby establishes a link between the SNR memory and aggression roles of the dCA2 region of the hippocampus as complementary processes, providing a broader understanding of how social memory regulates social behavior adaptively.

## Results

### Experience-dependent increase in aggression towards novel compared with familiarized conspecifics

To address whether and how social novelty relative to familiarity affects the levels of aggression, we designed a modified longitudinal version of the R-I test in which groups of residents (56 C57BL/6J male mice) encountered either the same initially novel intruder or a different novel intruder in one ten-minute session per day over five consecutive days (Fig. 1A). For each session, we measured the time the resident spent engaged in: 1. neutral social exploration of the intruder; 2. social dominance (e.g. mounting, chasing, intensive grooming of the intruder); and 3. aggression (defined as one or more episodes of biting attack of the intruder). We limited our analysis to the approximately 40% of residents that exhibited aggressive behavior on either the first or second day of the test (Fig. 1D). Importantly, the proportions of aggressive residents did not vary based on their randomly assigned group (Suppl. Fig. 1E).

**Figure 1:**
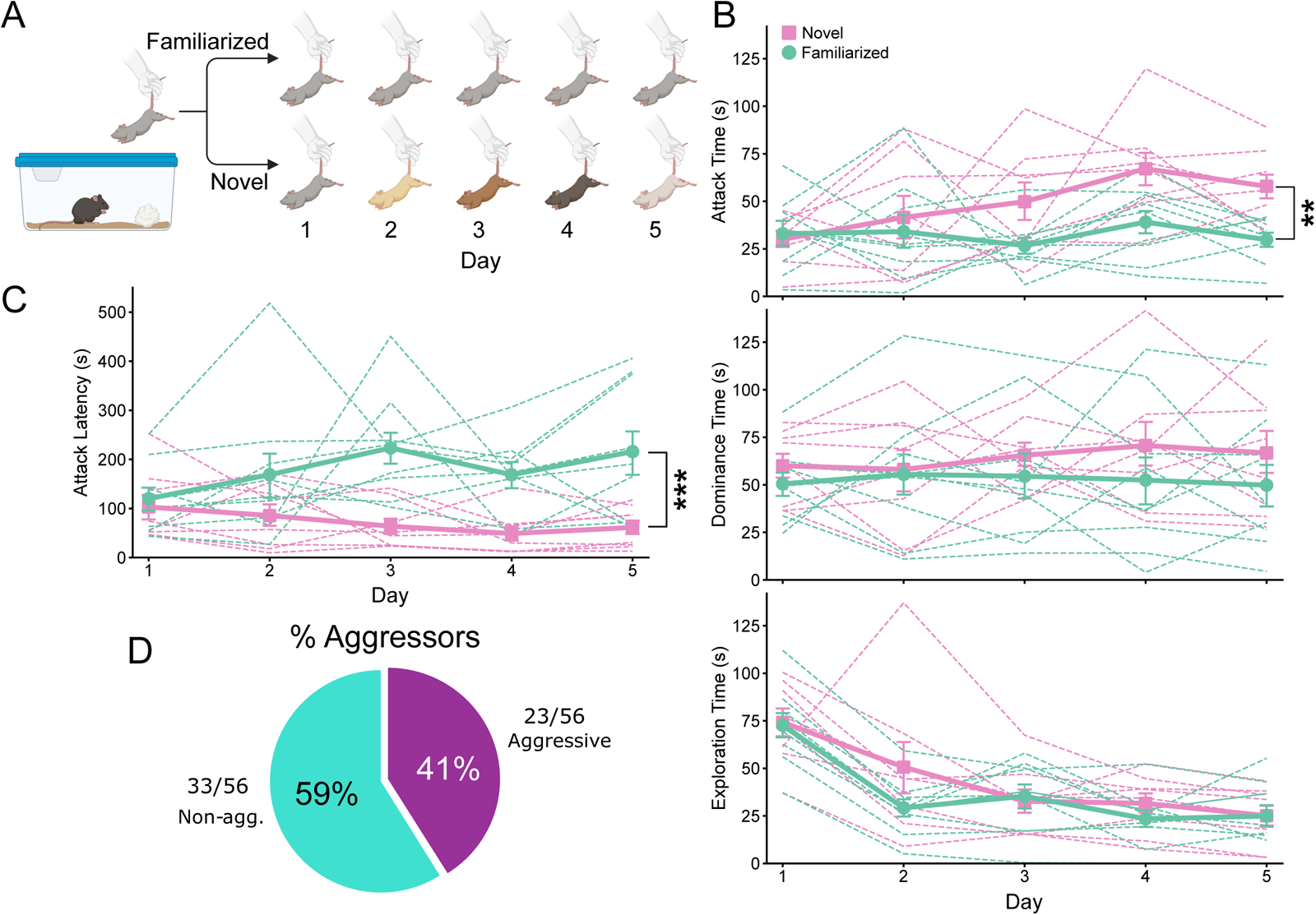
Experience-dependent increase in aggression towards novel compared to familiarized conspecifics in the longitudinal R-I test. **A**) Schematic and schedule of the 5-day modified resident intruder test for groups of resident mice in novel naïve and familiarized intruder conditions. **B**) Effect of intruder novelty on time spent in attack (top), dominance (middle), and exploration (bottom) with the same familiarized intruder on each day (green) or a novel intruder on each day (pink). Thick lines and error bars, mean ± SE. Thin lines, data for individual residents. Results from a Linear Mixed Model (LMM) show a significant effect of days on attack with the novel but not familiarized intruder (top panel). On Days 4 and 5, the group with the novel intruder (n = 8) exhibited significantly greater attack compared to the group with the familiarized intruder (n = 9; Day 4, *p* =0.026; Day 5, *p* = 0.026). No difference in dominance or exploration across groups and days. **C**) Latency to first attack for mice in groups with familiarized and novel intruders. There was a difference in days 3-5 between groups (LMM analysis), with day 3 and 5 reaching statistical significance (Day 3: *p* = 0.008; Day 4: *p* = 0.055; Day 5: *p* = 0.01). **D**) Proportion of mice that were aggressive across all groups and used in the analysis (no significant difference in proportions across each group—see S1E).

We found a significant difference in aggressive behaviors between the cohort that encountered a novel intruder on each day compared to the cohort that encountered the same intruder on each day, in both duration of attacks per session and latency to the first attack bout (Fig. 1B, top, Fig. 1C). Residents that encountered a novel intruder each day displayed escalating levels of aggression reported by both the total duration and latency to the first attack bout, as previously reported^9,11^. In contrast, mice that encountered the same, increasingly familiar intruder each day showed stable levels of aggression with no difference in duration or latency to first attack across days. By days 4 and 5, the differences in attack duration between the groups reached statistical significance. Additionally, time spent in attack was correlated with the number of biting episodes across the groups (Suppl. Fig. 1D). There was no significant difference between groups in time spent engaged in dominance or exploration behaviors during the five days of the test. Both groups of mice showed a significant decrease in social exploration after initial contact on day 1 (Fig. 1B, bottom). Although there was a trend for increasing time spent in dominance with the group that encountered novel intruders each day, this did not reach statistical significance (Fig. 1B, middle).

The increased aggression to the novel compared to familiarized intruders was specific to resident behavior and did not reflect the greater number of R-I sessions experienced by the intruder in the familiarized group (five sessions total) compared to the novel group (one session per intruder). Thus, when we controlled for the number of R-I sessions experienced by an intruder in a separate cohort of residents using intruders that were novel to the resident but that had undergone the same number of R-I sessions with other residents, we observed a similar increase in aggression by the resident to the experienced novel intruders over the five days of the test (Suppl. Fig. 1A-C).

These results suggest a resident-centric effect of social experience on aggression dependent on intruder novelty, with elevated levels of aggression to novel compared to familiar conspecifics. We examined this idea more explicitly in a second protocol in which the resident was exposed to the same, initially novel, intruder over 5 days, similar to the above protocol. However, on the sixth day, we introduced a second completely novel, but experienced intruder. Consistent with our first test, we observed stable aggression levels after day 3 across an initial 5-day R-I test towards a familiarized intruder. However, when confronted with a novel intruder on the following day 6, resident mice displayed significantly elevated aggression, with an increase in both duration and number of attacks and a lower latency to the first attack bout (Fig. 3C and Suppl. Fig. 3).

### Silencing of dCA2 attenuates aggression to novel, but not familiarized conspecifics

One hypothesis that could explain the increased aggression towards a novel intruder is that dCA2 pyramidal neurons, which enable SNR memory^25,26^, promote aggression towards a novel animal by providing a social novelty signal^24^, thereby enabling a mouse to discriminate a novel from familiarized intruder. To test this idea, we compared the effect of chemogenetic silencing dCA2 on aggression levels towards novel relative to familiarized intruders in an extended longitudinal R-I protocol (Fig. 2A, left). According to our hypothesis, we predicted that dCA2 silencing would preferentially decrease aggression towards novel compared to familiarized intruders.

**Figure 2:**
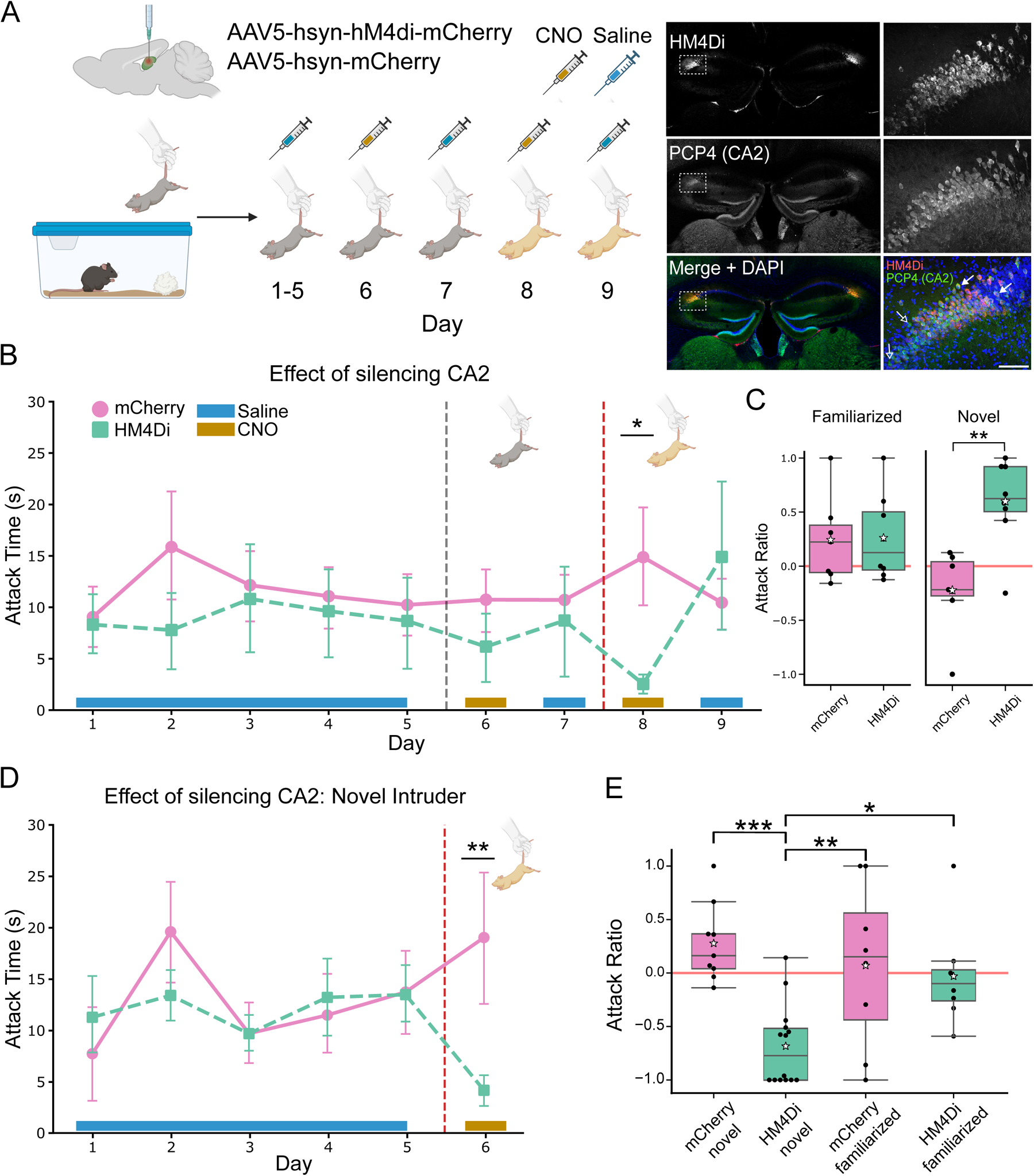
Silencing of dCA2 attenuates aggression to novel, but not familiarized conspecifics. **A)** Schematic and schedule for extended 9-day longitudinal R-I test. Mice were bilaterally injected in dCA2 with either hsyn-AAV5-hM4D(Gi)-mCherry (experimental) or hsyn-AAV5-mCherry (control) and R-I test begun 15 days later. Both groups were injected intraperitoneally with saline (days 1-5, 7, and 9) or CNO (days 6 and 8) 30 min prior to R-I test with either familiarized intruder (days 1-7) or novel intruder (days 8,9). Right Panel (left): coronal section showing dCA2 expression of hM4Di-mCherry (top), CA2 marker PCP4 (middle), and merged image co-stained with DAPI (bottom). Right Panel (right): 20x magnification of white bounded inset of pyramidal layer on left with corresponding staining. Scale bar 100 μm. Merged color image arrows: filled arrows indicate cells that are co-stained with PCP4 and mCherry, whereas empty arrows denote cells expressing PCP4 alone. **B)** Differential time spent in attack for mice in experimental and control conditions across 9 days. Experimental mice (n = 8, days 1-9) and control mice (n = 8, days 1-6; n = 7, days 7-9) injected with CNO on day 6 showed no differences in attack time to familiarized intruder (LMM fit). CNO significantly decreased attack time to a novel intruder (day 8) in experimental compared to control groups (LMM interaction effect, *p* = .022). **C)** Difference in attack time in absence and presence of CNO with familiarized (left panel: days 6 versus 7) and novel (right panel: days 9 versus 8) intruders for experimental and control groups expresses as attack ratio: (Attack time on day w/o CNO – Attack time on day CNO)/ (total attack time on both days). An independent two sample t-test revealed a significant difference between the groups with a novel (*t-stat* = 4.044, *p* = 0.00139) but not familiarized (*t-stat* = 0.088, *p* = 0.931) intruder. **D)** Effect of CA2 silencing on attack to novel intruder when performed on day 6. Experimental mice (n = 14) injected with CNO showed a significant attenuation of attack time compared to control group (n = 9) using a LMM (Day 6: *p* = 0.002). **E)** Difference in attack to novel and familiarized intruders when dCA2 is silenced on day 6 for experimental compared to control groups based on attack time ratio: (Day 6 attack time – Day 5 attack time)/ (Day 6 + Day 5 attack times). One-way ANOVA to compare the means, *F* (3, 35) = 8.428, *p* = 0.00024. Post hoc Tukey HSD test showed that the Experimental-Novel (Exp-Nov) group alone differed significantly from Control-Novel (Ctrl-Nov; *p* = 0.0003), Control-Familiarized (Ctrl-Fam; *p* = 0.0075) and Experimental-Familiarized (Exp-Fam; *p* = 0.024) groups, respectively.

As it was impractical to use the CA2-selective Amigo2-Cre mouse line to obtain sufficient aggressive animals, we used wild-type C57Bl/6J males and relied on the natural tropism of the AAV2/5 serotype to selectively infect dCA2 pyramidal neurons^24,30^. We thus bilaterally injected into dCA2 AAV2/5 expressing the mCherry-tagged hM4Di designer receptor exclusively activated by designer drugs (iDREADD, experimental group) or an AAV2/5 expressing mCherry alone (control group). Post-hoc staining with the CA2 marker PCP4 revealed near complete overlap with mCherry in dCA2 pyramidal neurons (Fig. 2A, right). All resident mice were injected with saline 30 min prior to the R-I test on days 1-5. Only mice that displayed one or more biting attacks within the first two days of the test were included in further analyses. Consequently, due to the variability in aggressive responses, the final number of mice retained for each experimental condition (control vs. inhibition) differed.

Three weeks following injection of iDREADD-or mCherry-expressing AAV, coinciding with day 6 of the R-I protocol with the familiarized intruder, the two cohorts were intraperitoneally injected with the iDREADD agonist CNO. After a 30-minute habituation, residents were exposed to the same familiarized intruder they encountered on days 1-5. Interestingly, we found no significant difference in the time spent in attack between groups, although there was a trend for decreased attack in the iDREADD group. We then repeated the R-I test following saline injection on day 7, a time when CNO from the previous day is no longer effective^3^, and observed normal levels of aggression in both groups of residents (Fig. 2C, left panel). Next, on day 8 of the test, we re-examined the effect of dCA2 silencing with CNO on aggression during an encounter with an experienced novel intruder. In keeping with earlier behavioral results, the control mCherry group exhibited greater time spent in attack towards the novel compared to the familiarized intruder (day 6-7). In stark contrast, the iDREADD group showed a marked decrease in attack behavior, to levels below that seen to the familiarized intruder, even when dCA2 was silenced. This decrease in aggression was reversible as aggression significantly increased to control levels following saline injection on day 9 in response to the same novel intruder presented on day 8 (Fig. 2B and C, right panel). What is of further interest is that the level of aggression was greater than that to the familiarized intruder the day after CNO injection (day 7), suggesting that the resident experienced the intruder on day 9 as being novel, despite having encountered it on day 8 when dCA2 was silenced. This result is consistent with previous findings that dCA2 is important for encoding familiarity and suggests that the resident failed to store a memory of the novel intruder when dCA2 was silenced.

In principle, the increased efficacy by which dCA2 silencing decreased aggression during the encounter with the novel compared to the familiarized intruder could be due to the difference in the order in which we tested the effects of CNO. To allow a more direct comparison of the results with familiarized compared to novel intruders, we repeated the R-I test in a different cohort of residents in which we first silenced dCA2 during presentation of the novel intruder on day 6, the same day that we had assessed the effect of silencing dCA2 during presentation of the familiarized intruder. Injection of CNO 30 minutes prior to the R-I test caused an almost complete suppression of attack behavior towards the novel intruder, which was a much greater effect than seen with dCA2 silencing when a familiarized intruder was presented on day 6. In contrast, the control mCherry group of residents showed the expected increase in attack to the novel intruder compared to the familiarized intruder (Fig. 2D). We quantified the effects of dCA2 silencing by measuring an attack ratio—the total duration of attack on day 5 (saline injection) minus the attack duration on day 6 (saline injection) divided by the sum of attack durations on the two days. We observed a significant decrease in attack time to novel intruders in the dCA2-silenced cohort relative to the control cohort. The attack time in the dCA2-silenced cohort presented with a novel intruder was also significantly less than the attack time in both the dCA2-silenced and control cohorts presented with a familiarized intruder (Fig. 2E, Suppl. Fig 2C).

Thus, our chemogenetic silencing results confirm previous findings that dCA2 promotes aggression^22^ and show that this effect is more pronounced with novel compared to familiarized intruders. These results are consistent with the hypothesis that dCA2 may control aggression through its role in SNR memory, informing downstream brain regions as to whether an intruder is novel or familiar, and promoting aggression accordingly. Interestingly, whereas silencing dCA2 decreased aggression to the novel intruder on day 6, the time spent in exploration towards the intruder significantly increased, with no change in dominance, suggesting a shift in the probability of behavioral responses (Suppl. Fig 2A, B).

### Single cell activity in dCA2 differentially responds to novel and familiar intruders

In addition to signaling social novelty, dCA2 might also promote aggression by encoding the distinct behavioral states of social exploration, dominance, and aggression present in the R-I test. This view is consistent with prior fiber photometry results showing progressively larger increases in mean calcium signals in the population of dCA2 pyramidal neurons during episodes of social exploration, dominance and attack^22^. However, whether these neural responses are modulated by social novelty and how they are produced at the single cell level is unknown. We therefore examined whether the dCA2-dependent enhancement in aggression towards novel intruders was associated with differences in the representations of social interactions with novel compared to familiar intruders.

To explore these ideas, we expressed the genetically encoded calcium indicators GCaMP6f or GCaMP8s in dCA2 pyramidal neurons using injections of AAV2/5 serotype in dCA2. Post-hoc staining with the CA2 marker PCP4 revealed strong overlap with GCaMP expression in dCA2 pyramidal neurons (Fig. 3B). We performed microendoscopic calcium imaging 5-6 weeks after viral injection to monitor dCA2 pyramidal neuron activity in freely moving mice. Residents underwent the longitudinal R-I test beginning with presentation of a novel intruder on day 1, followed by five consecutive days with the same intruder. Finally, on the sixth day, residents again encountered a novel intruder (Fig. 3A).

**Figure 3:**
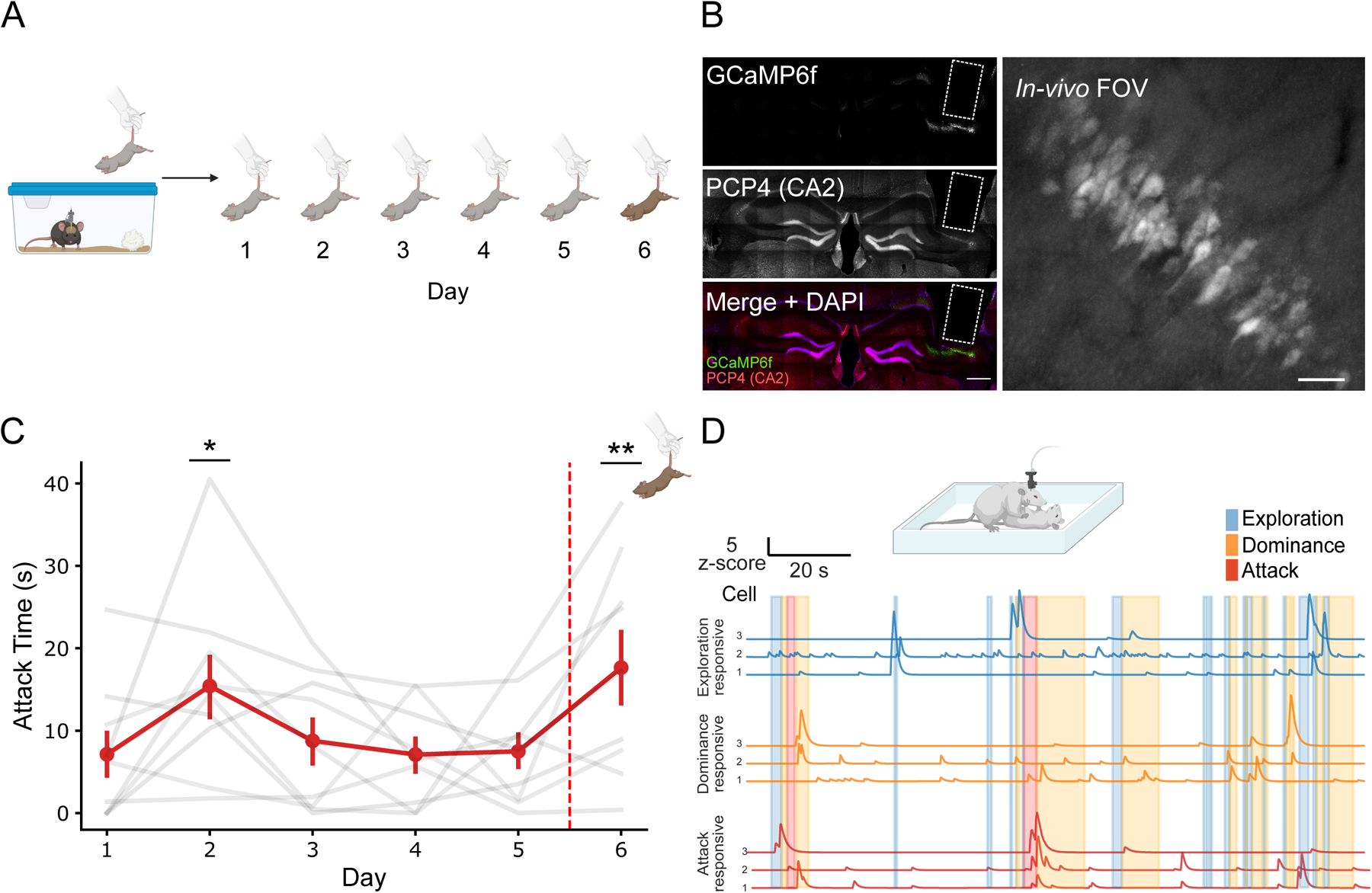
Imaging dCA2 activity during social aggression to novel and familiarized conspecifics. **A)** Schematic and schedule for imaging dCA2 calcium signals across 6-day longitudinal sessions. hsyn-AAV 2/5 serotype mediated targeting of dCA2 was used to deliver GCaMP6f or GCaMP8s. A 10° angled GRIN lens was implanted on top of the alveus and positioned for imaging over dCA2 with a docking baseplate for multi day recording. **B)** Post hoc confocal image of a brain section from an example mouse. Counter-staining with the CA2 marker PCP4 (middle) confirms the expression of GcaMP6f (top) in CA2 PNs. Scale bar 500 μm. Right, in vivo one-photon image (maximum intensity projection) from a representative field of view showing GcaMP6f fluorescence in cell somata and dendrites of dCA2 PNs. Scale bar 100 μm. **C)** Time spent in attack for all imaged aggressive mice across days (n = 9). Presentation of an experienced novel intruder increased the time spent in attack on days 2 and 6 (LMM, p-values (p <.05), Day 2: 0.025, Day 6: 0.004). **D)** Synchronized calcium signals and behaviors. Normalized and background subtracted calcium signals for three example cells activated during manually annotated bouts of exploration (blue), dominance (orange), or attack (red) behaviors from one resident during an R-I session.

Despite the presence of the gradient refractive index (GRIN) lens implant and attached cable, residents engaged in typical dominance behaviors during interactions with the initially novel mouse during familiarization on days 1–5 and during presentation of a second novel intruder (that had previously undergone five days of attack by a different resident) on day 6 (Suppl. Fig. 3). In this set of experiments, we found a significant increase in attack behavior on day 2 compared to day 1 with the same intruder, presumably due to enhanced experience of the resident in attack behavior^9,15,16^. Residents then displayed stable, reduced aggression levels on days 3,4, and 5 to the now familiarized intruder, presumably because the resident encoded the reduced perceived threat posed by the familiarized intruder (Fig. 3C). On day 6, upon presentation of an experienced novel intruder, resident mice displayed a significantly elevated total duration of attack behavior, an increase in the number of attacks, and a lower latency to the first attack bout (Fig. 3C and Suppl. Fig. 3D). Additionally, on days 3,4, and 5, we observed significant differences in exploration and dominance behaviors towards the familiarized intruder compared to day 1 (Suppl. Fig 3 A-B).

We next asked whether and how dCA2 activity encodes different social behaviors, how this may change over the course of 5-day presentation of the same intruder, and whether there are differences in representations of behaviors during the presentation of the novel intruder on day 6. We performed frame-by-frame annotation of social behaviors (exploration, dominance, and attack) from synchronized neural activity and video recordings (refer to Table 1 in the methods section for operation definitions). Annotations were then aligned to post-processed calcium signals, resulting in denoised single cell calcium transients^31,32^ suitable for downstream analysis (Fig. 3D, see Methods).

We first investigated single-cell response profiles by comparing activity during social exploration, dominance, or attack with baseline levels of activity, defined as periods when the resident was not interacting with the intruder. Neuron responses were deemed activated or inhibited by a given behavior if the mean difference in activity with baseline was significantly more positive (activated) or negative (inhibited) than chance, defined as a difference that was greater or less than 95% of a null distribution consisting of 1000 random assignments of the calcium traces to a behavior label (α = 0.05). A neuron whose activity did not exceed the thresholds for any behavior was considered non-responsive.

Based on an analysis of an average of 586 neurons per day from a pool of 9 aggressive mice, we found a significant proportion of cells that were either activated or inhibited during one of the three annotated behaviors compared to the non-social interaction baseline (Fig. 4A, see Fig. 3D for examples of activated response cells). Averaged across the six days of recordings without tracking individual cells across sessions, we found that 24.4%, 31.4%, and 12.5% of cells were activated during exploration, dominance, and attack respectively (compared to chance levels of 5%). A somewhat smaller fraction of cells was inhibited during the behaviors, with 17.3% inhibited during exploration, 15.9% during dominance and 5.8% during attack. Although at first glance we did not see striking changes across the days, a statistical analysis revealed a number of significant differences.

**Figure 4:**
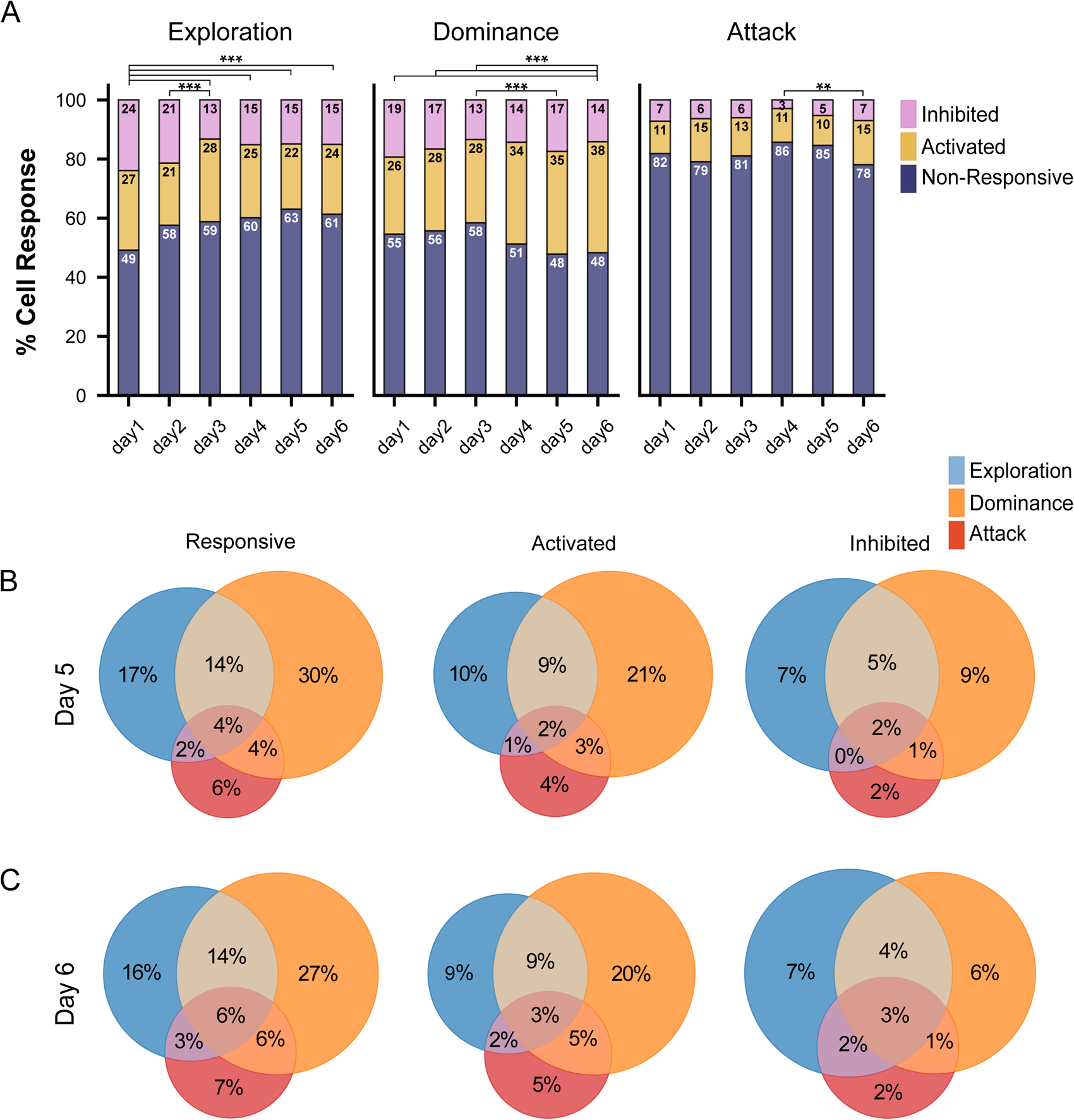
Social behavior encoding in resident dCA2 single cells during encounters with familiarized or novel intruder. **A)** Percent of dCA2 cells whose activity was increased (activated, yellow), decreased (inhibited, pink) or unchanged (unresponsive, blue) during bouts of exploration, dominance, or attack behaviors compared to baseline activity during non-social episodes. Activated and inhibited cells responses judged significant if magnitude of change was, respectively, greater or less than 97.5% of responses in 1000 shuffled data sets (n = 6, 9, 8, 7, 8, 9 attacking resident mice during encounters with increasingly familiar intruder on days 1-5 and novel intruder on day 6, respectively). Total cells = 502, 646, 619, 482, 625, 644 for days 1-6, respectively. Cell response distribution during attack differed between day 6 compared to day 4 (χ2 statistic: 13.23, *p* = 0.0013; alpha = (0.05)/15 = 0.0033; χ2 statistics with Bonferroni correction for multiple comparisons). Cell response distributions during exploration differed between day 1 and days 3, 4, 5 and 6 and day 2 compared to day 3. Cell response distributions during dominance differed between day 6 and days 1,2 and 3 and between days 3 and 5. **B-C) Left,** Percent of cells responsive (activated or inhibited) to one or more of the three social behaviors; Middle, percent of cells activated by one or more of the three behaviors; **Right**, percent of cells inhibited by one or more of the three behaviors. Measurements from day 5 (B) or day 6 (C). Chi-square test of independence for responsive cells (combined fraction of activated and inhibited cells) revealed no significant difference above chance in proportion of cells responsive to two behaviors at *p* = 0.05 after Bonferroni correction for multiple comparisons (alpha = (0.05)/3 = 0.016). Percent of cells activated during both exploration and dominance (but not during attack and either exploration or dominance) was greater than chance for both days (day 5, *p* = 0.0065; day 6, *p* = 0.0015). For inhibited cell populations, all binary comparison differences were greater than chance. *p<0.05, **p<0.01, ***p<.001

During the initial five days of interaction with the increasingly familiar intruder, there was a marked decrease in the fraction of cells that were inhibited during social exploration behavior, dropping from 24% of cells on day 1 to only 13% by day 3. The difference between day 1 was significant on days 3-6 and between day 2 and day 3 (chi-squared test of independence, p-value adjusted for 15 pairwise comparisons between days using a Bonferroni correction). In contrast, the proportion of activated cells during social exploration remained stable throughout the six-day period, with no significant variations detected (Fig. 4A, left).

For dominance related behaviors, there was a significant increase in the proportion of activated cells from days 1-3 (∼27%) to day 6 (38%) and between days 3 and 5, with no apparent differences in the fraction of inhibited cells (Fig. 4A, middle). Cell responses during attacks also evolved over the days, culminating in a significant increase in both activated (from 11% to 15%) and inhibited cell responses (from 3% to 7%) between days 4 (familiar intruder) and 6 (novel intruder), with a similar trend that did not reach significance between days 5 (familiar intruder) and 6 (Fig. 4A, right). Overall, these results suggest a subtle change in dCA2 neural activity associated with the evolution of resident behavior during familiarization with the same intruder on days 1-5 and during the encounter with a novel intruder on day 6.

We next investigated whether the responsive neurons were specialized for a specific behavior or whether the same cell could respond to multiple behaviors by plotting a Venn diagram charting the percent of responsive cells (either activated or inhibited) in the seven possible categories of overlapping responses for the three behaviors (Fig. 4B). We then asked whether the fraction of cells that responded (either activated or inhibited) to more than one behavior was greater or less than that expected by chance overlap. For example, on day 5 (Fig. 4B, left), approximately 37% of cells responded during social exploration, 52% of cells responded during dominance, and 16% responded during attack. The 18% of cells that responded to both exploration and dominance behaviors was not significantly different than the approximately 20% overlap expected by chance (percent overlap assuming statistical independence = 0.37 x 0.52 x 100% = 20%; p = 0.45; chi square test of independence). Similarly, the 4% of cells that responded to all three behaviors was not different than expected by chance overlap (0.37 x 0.52 x 0.16 x 100% = 3.1%; p = 0.29). A similar conclusion was reached when analyzing responsive cell data from day 6.

The same analysis of response overlap was then performed separately on activated and inhibited response cell categories (Fig. 4B-C middle, right). We found that for the inhibited cells on both days each binary behavior comparison was significantly greater than expected from chance overlap (Fig. 4B-C right). For the activated cell category on day 5, the overlap between cells activated by both dominance and exploration was significantly greater than chance, while attack response overlap with exploration and dominance did not differ from chance (Fig. 4B-C right). On day 6, there was again a significantly greater overlap in cells responding to both exploration and dominance compared with chance, with a significant overlap in attack and dominance responses (p = 0.027) and a trend for increased overlap in attack and exploration responses (p = 0.074) and over chance levels (Fig. 4B-C middle).

To test the hypothesis that dCA2 responds more vigorously to a novel intruder compared to a familiarized one, we focused our analysis on cell responses from days 4 and 5 (when exposed to a fully familiarized intruder) and compared them to responses on day 6 (when exposed to a novel intruder). We first compared the different types of cell responses (activated, inhibited, and non-responsive) for each behavior across the three days. We found a significant increase in the proportion of cells responding to attack from days 4 to 6 (Z-statistic: -3.23, p-value: 0.0012) and from days 5 to 6 (Z-statistic: -2.99, p-value: 0.0028). When analyzing each behavior by response type (activated or inhibited), we found a significant increase in the fraction of inhibited neurons from day 4 to 6 (χ² (2) = 8.45, p = 0.0036) and activated neurons from day 5 to 6 (χ² (2) = 6.31, p = 0.012). Moreover, the fraction of cells responsive during exploration or dominance behaviors remained consistent across these days.

To control for the possibility that the differences in cell responses between days 5 and 6 was due to the difference in number of R-I sessions experienced by the resident and not the difference in familiarity of the intruder, we performed the same analysis comparing days 4 and 5 with the same familiarized intruder. We found no significant difference in responsive cells for any behavior category between these days (χ² (2) = 4.1, p = 0.13). Further, no significant difference was found when activated and inhibited neuron categories were separately tested. Our findings of a significant difference in the fraction of responsive cells during attack of a novel intruder on day 6 compared to attack of the familiarized intruder on days 4 or 5, but no difference between days 4 and 5, is consistent with our behavioral results demonstrating a large increase in attack duration and frequency towards the novel intruder (Fig. 3D).

### Dynamics of dCA2 neuron calcium responses at onset of resident behaviors

Our finding that a lesser proportion of cells were activated during resident attacks compared to social exploration or dominance was unexpected, especially considering the significantly larger increase in attack-specific behaviors towards a novel intruder on day 6. However, the simple measure of cell proportions fails to provide information as to the dynamics and magnitude of dCA2 cell responses at the onset of different behaviors. We therefore aligned the activity of each cell to the start of a given behavior and determined the average responses of each cell for each behavior on a given day. Cells were then tracked over several days^31^. We focused on responses during days 4,5, and 6 to compare activity levels of the same putative cells with the same intruder (days 4 versus 5) or with different intruders (days 4 or 5 versus 6).

We separately determined the average behavior-aligned response magnitudes of the subpopulations of activated and inhibited neurons identified in Figure 4 for each of the three behaviors (Fig. 5). The subpopulation of activated cells showed a clear, time-dependent increase in the mean calcium signal aligned to the start of both exploration and dominance behaviors of both the familiarized intruder (day 5) and the novel intruder (day 6) (Fig. 5A). For both days, the mean calcium responses of the population of cells that were activated during exploration and dominance behaviors was significantly greater 0-1 s after the onset of a behavior (Fig. 5B, top) and compared to a baseline 2-3 s prior to behavior onset, analyzed through absolute difference (Fig. 5B, bottom). In addition, calcium transients for attack-activated cells demonstrated a robust and significant time-locked response on day 6, with a mean increase significantly greater than that observed for the dominance or exploration responsive cells, both when measured after behavior onset and absolute z-scored difference from baseline (Fig. 5B, right).

**Figure 5:**
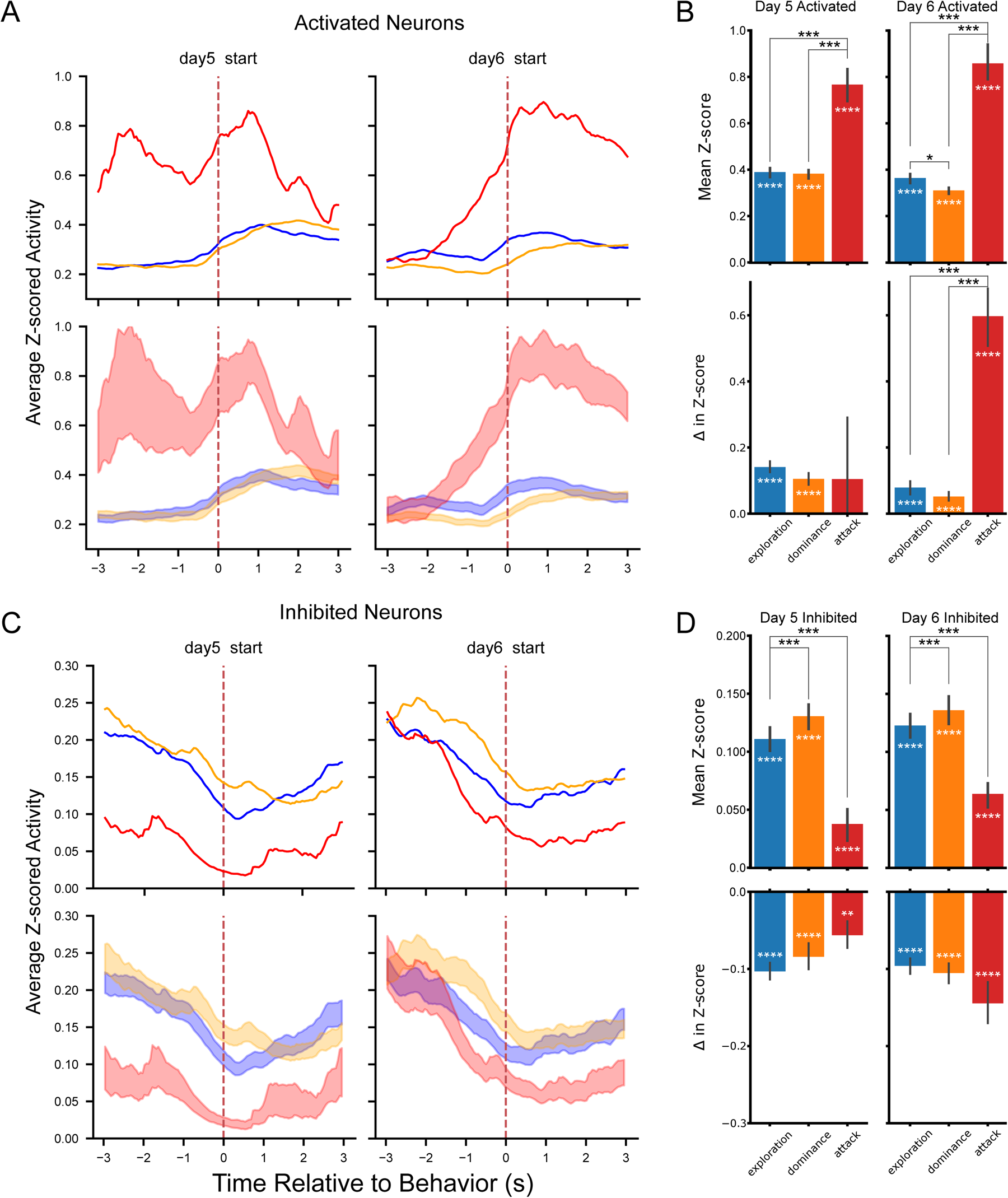
dCA2 neuron response dynamics during social encounters with familiar and novel intruders. **A)** Mean *z-scored* responses (top) and SE (bottom) of behavior activated neurons aligned to onset (dashed line) of attack bouts (red), dominance bouts (orange) and exploration bouts (blue) during interaction with familiarized (day 5) and novel (day 6) intruder. Cell counts for days 5 and 6, respectively: Exploration = 138, 152; Dominance = 217, 242; Attack = 63, 96. **B)** The bar graphs show the mean *z-scored* calcium signal of neurons activated for respective behaviors on days 5 and 6. **Top**, mean ± SEM (**top**) 0-1 s after behavior onset; **Bottom**, difference in z-scored activity (z-score averaged 0-1 s after behavior onset minus z-score averaged 2-3 s before behavior). Significance for each bar was determined against a null hypothesis of zero difference using a one-sided t-test. Comparisons between each behavior within a day made using independent t-test. **C)** Similar to Panel A but for inhibited neurons. The graph shows the mean z-scored responses (top) and SE (bottom) aligned to behavior onset (dashed line). Cell counts for days 5 and 6, respectively: Exploration: 93, 97; Dominance: 109, 91; Attack: 33, 45. **D)** Similar to Panel B showing z-score responses for inhibited neurons. The same statistical test was applied as in Panel B. *p<0.05, **p<0.01, ***p<.001.

The dynamics of the calcium responses of attack-activated neurons on day 5 were more complex, with a significant elevation in the baseline periods of non-attack behaviors even 3 s prior to the onset of attack. Nonetheless, there was a significant increase in calcium levels 0-1 s after attack onset but not in absolute z-score difference from baseline (Fig. 5B, left). Despite the complexities of the day 5 dynamics, the calcium response on day 6 relative to baseline during attack of the novel intruder was significantly greater than that on day 5 during attack of the familiarized intruder. In contrast, the calcium responses during exploration and dominance on day 6 were significancy decreased than those during the same behaviors on day 5 as a measure of absolute difference (Fig. 5B, bottom).

We adopted the same approach to analyze the dynamics of the responses of neurons that were inhibited during a specific behavior on days 5 and 6. There was a significant decrease in calcium activity aligned to the onset of the three behaviors on both days compared to baseline (Fig. 5C). In addition, for both days we found a significant difference in the mean z-scored activity 0-1 s after behavior onset of attack compared to dominance and exploration (Fig. 5D, top). There was a trend for an increasing magnitude of inhibition during exploration compared to dominance compared to attack behaviors on day 6, while we observed an opposite decreasing trend for the calcium responses aligned to the three behaviors on day 5 (Fig. 5D, bottom). Furthermore, we found a significantly larger inhibitory response magnitude aligned to attack behavior on day 6 compared to day 5, with no difference for exploration- or dominance-aligned inhibitory responses (Fig. 5D, bottom). Overall, these results demonstrate a consistently larger attack-aligned calcium responses compared to baseline during interactions with the novel intruder on day 6 compared to the familiarized intruder on day 5 for both activated and inhibited cells. Interestingly, exploration- and dominance-aligned responses were significantly greater during interactions with the familiarized compared to novel intruder for activated cells, but no differences were observed for inhibited cells.

One potential confounding factor in the calcium activity aligned averages is that the baseline measured 3 to 2 s prior to a behavior may depend on the particular behavior in which a given mouse was engaged at this time. For example, for transitions to attack behavior the animal could have been engaged in non-social exploration, social exploration, or dominance prior to attack. As a result, neuronal responses and their preceding baseline activity aligned to a given behavior may differ across days if the probability of engagement in a particular behavior preceding the transition were to vary. However, we found no difference in the sequence of behavioral transitions to an attack bout across days 4, 5, or 6 (Suppl. Fig. 4C).

We next examined whether single neuron responses on the different days for a given behavior were correlated. We further tested whether any such correlations differed during interactions with the same familiarized intruder on successive days from the correlation in activity during interactions across days with the familiarized compared to novel intruder. We constructed population activity vectors by averaging the activity of each cell aligned to the onset of each behavior on each day bounded by a 500-ms window. For display purposes we sorted the single cell responses by their mean activity on day 6 (Fig. 6A, see Suppl. Fig. 4A for alignment based on day 5). Next, we tested for any significant correlation in the population activity between days in a pairwise manner using Spearman’s rank-order correlation and a Bonferroni correction for multiple comparisons. This approach revealed a significant correlation in dCA2 activity between each pair of days (days 4-5, 4-6, and 5-6) during exploration and dominance behaviors. However, the magnitude of the correlation was greater for days 4 to 5 on average compared to days 5 to 6 or days 4 to 6. We also found a significant correlation between activity during attack behaviors with the same familiarized intruder on days 4 to 5. In contrast, there was no significant correlation between the activity recorded during attack of the novel intruder on day 6 compared to the familiarized intruder on either day 4 or 5. These results are summarized in the confusion matrix of Figure 6B. To assess further the significance of the observed correlations, we calculated the correlations with randomly shuffled cell identities across pairs of days for each behavior. Comparing our observed correlation with the null distribution confirmed our initial findings (Suppl. Fig. 4B). We thus conclude that individual dCA2 neurons show significant correlations in their responses to the same behaviors across days, but that the correlations in cell activity were greater across sessions with the same animal compared to sessions with different animals.

**Figure 6:**
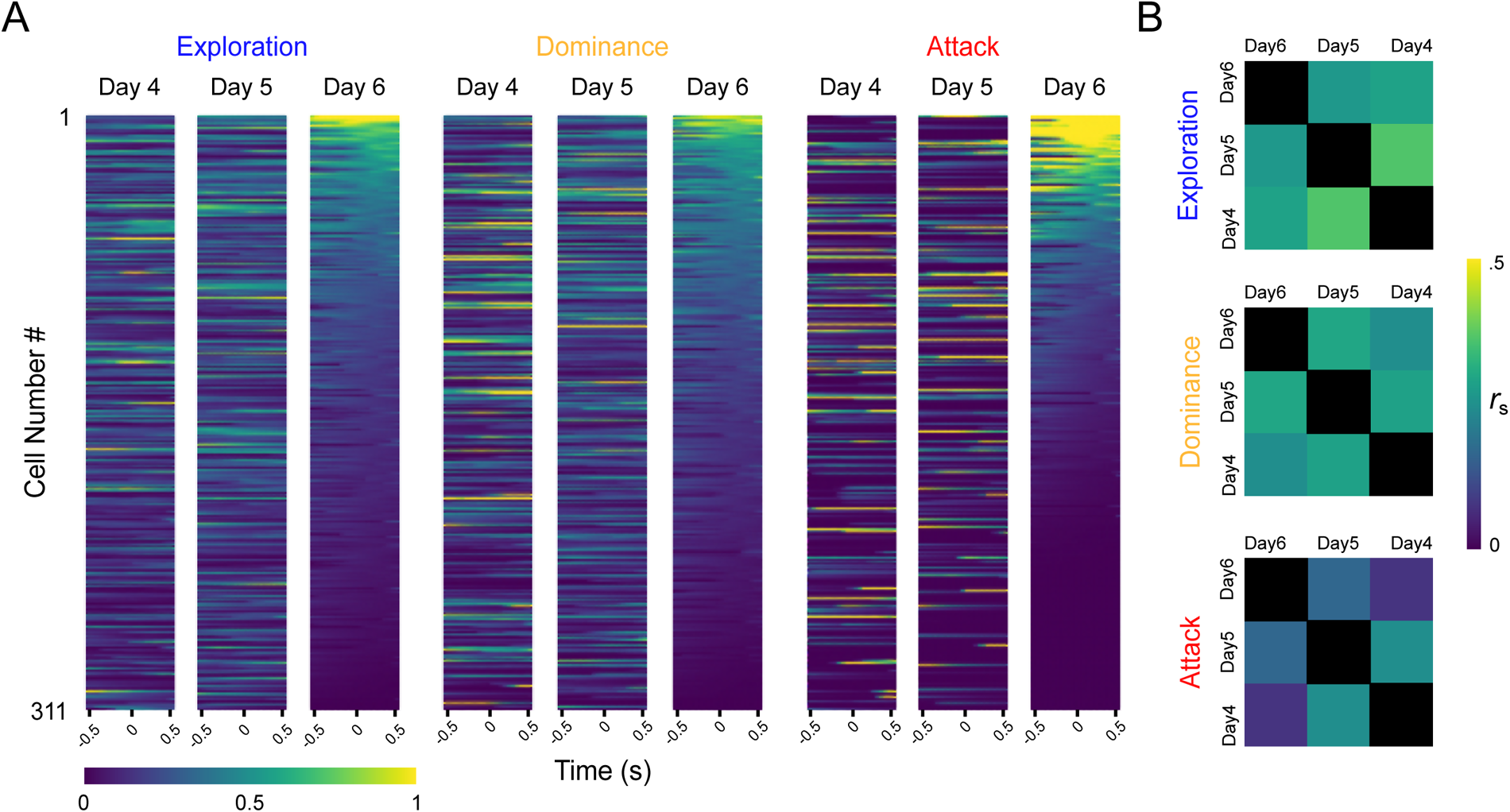
dCA2 activity correlation across days. **A)** Average neural activity vectors for 311 tracked neurons from n = 6 aggressive mice on days 4, 5, and 6 aligned to behavior onset bounded by 500 ms windows. Neurons are corresponding across rows and sorted based on day 6 mean activity measured 0 to 500 ms after behavior onset. **B)** Correlation matrix for pairwise comparison of days calculated based on mean activity correlation from 0 to 500 ms after behavior onset. Correlation reported by spearman’s rank and significance by alpha = (0.05)/3 = 0.0167, Bonferroni correction for multiple comparisons. Day 6 and 5; attack: *r_s_* = .166, *p* = 0.137, exploration: *r_s_* = 0.266, *p* = 6.14e-06, dominance: *r_s_* = 0.296, *p* = 3.62e-07; Day 6 and 4; attack: *r_s_* = .076, *p* = 1, exploration: *r_s_* = 0.288, *p* = 7.71e-07, dominance: *r_s_* = 0.244, p-value = 4.36e-05; Day 5 and 4; attack: *r_s_* = 0.254, *p* = 0.009, exploration: *r_s_* = 0.363, *p* = 1.34e-10, dominance: *r_s_* = 0.285, *p* = 1.04e-06.

### Assessing the accuracy of population-based decoding of social behaviors with novel compared to familiar intruders

The above single cell analysis depends on arbitrary activity thresholds for response classifications and does not take into account the potential importance of weakly-selective or mixed-selectivity neurons or cell correlations. We thus next determined whether and how the population activity of dCA2 pyramidal neurons encodes exploration, dominance, and attack behaviors during interactions with novel and familiar animals using a linear classifier to decode the three possible pairs of behaviors. We trained a support vector machine (SVM) classifier with a linear kernel to distinguish between manually annotated behavior dichotomies (exploration versus dominance, exploration versus attack, and dominance versus attack) based on the aligned calcium signals for each of the subject mice examined on each of the six days of the resident-intruder test (Fig. 7A). We then tested the accuracy of the classifier on decoding behaviors for a subset of calcium activity data not used for training. Because the mice engaged in behaviors for differing amounts of time, we subsampled the data to balance the time spent in each behavior for a given dichotomy for a given subject mouse. We calculated chance levels of decoding by random shuffling of the calcium signals relative to the behaviors.

**Figure 7:**
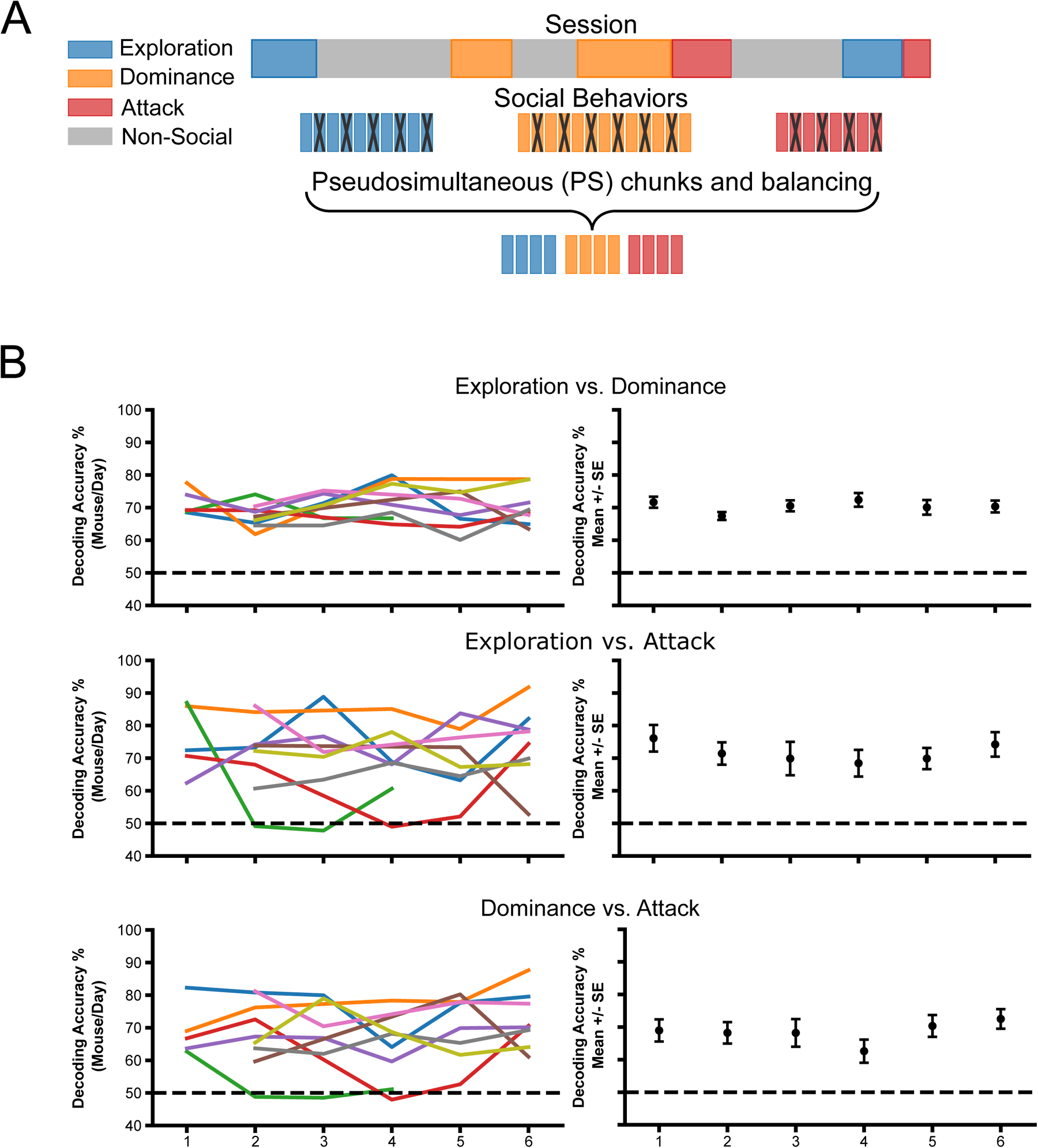
Decoding social behaviors during interactions with familiar and novel intruder. **A)** Schematic of decoding approach. Periods of a given social behavior were extracted from full sessions and concatenated to form behavior vectors that were then divided into 500 ms bins of non-contiguous pseudo simultaneous (PS) trials to avoid autocorrelation artifacts. Calcium activity data for binary comparison classes were balanced and split into training and testing data sets for SVM decoding. **B) Left,** Single resident mouse decoding accuracies across days during R-I test for indicated behavior dichotomies using all available neurons for each mouse on each day. **Right,** mean +/-SEM decoding accuracy for data from all aggressive resident mice across days (n= 5, 9, 6, 7, 8, and 8 aggressive mice and 455, 646, 307, 420, 482, 625, 605 neurons on days 1,2,3,4,5, and 6, respectively). There was no significant difference in decoding accuracy across days.

We found that separate classifiers trained on the different dichotomies accurately predicted behavior classes significantly above chance levels on each of the days of the test (Fig. 7B), with no obvious trends for changes in decoding performance across days. Although decoding of exploration versus dominance appeared to be more consistent across days and subject mice compared to the decoding of attack versus dominance and attack versus exploration, the data are not strictly comparable due to differences in the time spent engaged in the different behaviors across the different resident animals and days. This is particularly so with attack behavior, which was much sparser than exploration or dominance and varied considerably among days and resident mice. Moreover, although we imaged the same region of dCA2 on each day, we did not compare the activity of the same neurons across the different days. These factors, together with differences in the number of neurons recorded from day to day and across residents over the six days of the recordings, limit the ability to draw inferences from comparisons in decoding performance across the days of the experiment.

Thus, to enable comparisons across days and animals, we longitudinally registered the same neurons^31^ in a given animal over triplets of successive days. We confined the registration to three-day periods because of dropout of sequentially registered neurons over longer time frames. Moreover, because of differences in durations of different behaviors, we balanced the data to analyze neural activity in identical-sized blocks during the three behaviors across days and animals, which enabled us to directly compare SVM classifier accuracies. We tracked neurons over two triplets of days: A. Days 3,4,5, and B. Days 4,5,6. The first triplet corresponds to a time-period during interactions with the same highly familiarized intruder when aggression levels had stabilized (see Fig. 3C and Suppl. Fig. 3). The second triplet allowed us to compare the responses of the same cells during interactions with the familiarized compared to novel intruder. To account for different times spent in behaviors across triplets of days, we combined training data on days 4 and 5 to match the increased time spent in attack with the novel mouse on day 6. We then tested decoder performance on either familiarized day 3 (n = 6; Fig. 8B) or on novel day 6 (n = 8; Fig. 8A).

**Figure 8:**
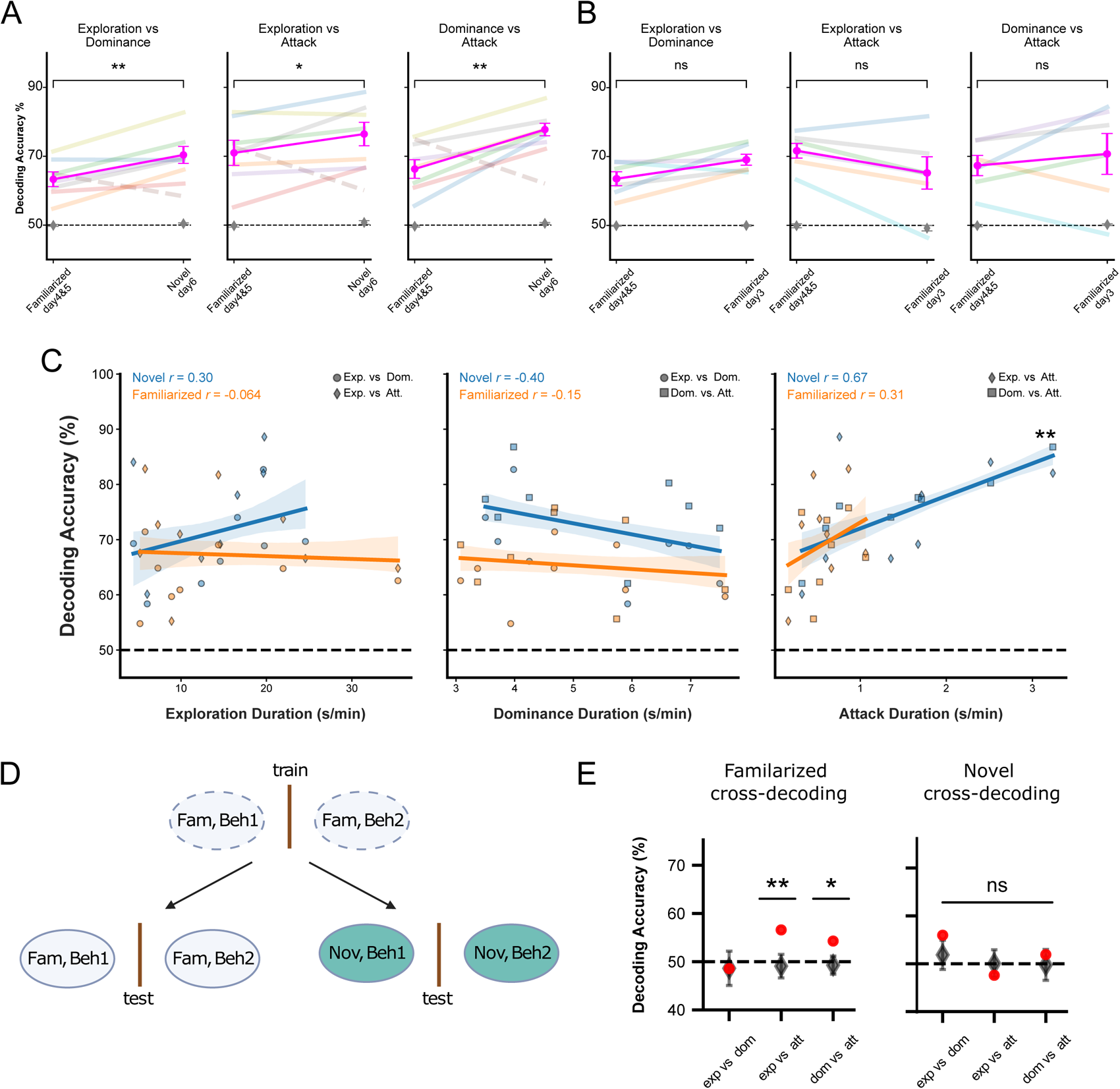
Comparison of behavioral decoding accuracy during R-I test with familiar and novel intruders. Calcium activity was measured from the same set of neurons tracked across triplets of days with the same familiar intruder (days 3,4,5) or across triplets of days including the familiar and novel intruder (days 4,5, and 6). Data were balanced to obtain similar interaction times for all binary behavior comparisons. **A)** Decoding accuracy of behaviors was significantly greater during interactions with the novel intruder on day 6 compared to the interactions with the familiarized intruder using combined data from days 4 and 5. Exploration versus dominance: *t-stat =* 4.24, *p =* 0.005; exploration versus attack: *t-stat* = 2.77, *p =* 0.032; and dominance versus attack: *t-stat* = 5.91, *p =* 0.00104 (paired t-tests; n = 7; data from one aberrant mouse that showed decreased attack to the novel versus familiarized intruder was excluded from analysis). **B)** No significant difference in decoding accuracies during interaction with familiar intruder on day 3 compared to combined data with same intruder on days 4 and 5. Exploration versus dominance: *t-stat* = 2.22, *p* = 0.077; exploration versus attack: *t-stat =* 2.32, *p=* 0.067; and dominance versus attack: *t-stat =* 0.79, *p =* 0.46 (paired t-tests; n = 6). **C)** Decoding accuracy for indicated behavior dichotomies plotted against duration (seconds, s) a resident was engaged in exploration (left), dominance (middle) or attack (right) of familiarized (orange) or novel (blue) intruders, normalized by the total time spent in the arena (minutes, min) across days of the R-I test. Pearson’s correlation coefficient is reported. Exploration versus dominance or attack: Novel intruder, *r* = 0.30, *p =* 0.26; familiarized intruder, *r* = -0.064; *p =* 0.814); Dominance versus attack or exploration: Novel intruder, *r* = -0.40, *p =* 0.126); familiarized intruder, *r* = -0.15, *p =* 0.573); Attack versus exploration or dominance: Novel intruder, *r* = 0.67, *p =* 0.005; familiarized intruder, *r* = 0.31, *p =* 0.24. **D)** Cross-day decoding of behavioral dichotomies from same tracked neurons. (Left) Schema of training classifier to decode behavioral dichotomies based on data from either day 4 or 5 (chosen based on maximal attack time per mouse) and tested on data from either familiarized day 3 or novel day 6. The reverse training/testing scheme was implemented to obtain a mean score. **E**) **Left**, Performance of linear classifier for decoding behavioral dichotomies with same familiarized intruder on training and testing days. Exploration versus dominance decoding accuracy = 0.485; *p =* 0.491; exploration versus attack accuracy = 0.565, *p =* 0.001; dominance versus attack accuracy = 0.542, *p* = 0.0039. **Right**, Decoding performance with data from a familiar intruder used for training and novel intruder used for testing (and vice versa). Exploration versus dominance accuracy = 0.559, *p* = 0.089; exploration versus attack accuracy = 0.475, *p* = 0.194; dominance versus attack accuracy = 0.519, *p* = 0.251). Red circles show the average decoding performance from 20 cross-validations. Black circles and error bars show mean ± 2 SD of shuffled data. P-values are given by z-score of performance values compared to the null model distributions (one sided t-test). Decoding performances from n = 399 neurons and 8 mice.

Decoding performance across the triplet of days during interaction with the same familiarized intruder (triplet A) was significantly above chance for all three dichotomies and did not differ significantly across days. In contrast, when we compared decoding performance across the triplet of days during interactions with the familiarized compared to novel intruder (triplet B), we found an increase in decoding performance in 8 out of the 9 animals we examined for each of the three behavioral dichotomies during interactions with the novel compared to familiarized intruder. Interestingly, the one resident mouse that decreased its attack time with the novel compared to familiarized intruder showed a large decrease in decoding accuracy with the novel compared to familiarized intruder. When we excluded this aberrant resident from the analysis, we found a significant increase in decoding accuracy with the novel intruder for all three dichotomies compared to the decoding accuracy the familiarized intruder (Fig. 8A). To rule out any potential confounds in combining attack behavior during days 4 and 5, we also analyzed our results by considering days 4 and 5 separately, selecting the day with maximal attack time and passing a minimal number of bouts of attack for each resident. We again found an enhanced decoding of interactions with novel compared to familiar intruders when we excluded the aberrant mouse (Suppl. Fig. 5B, D). Even with the inclusion of data from the aberrant resident, decoding accuracy increased significantly towards novel compared to familiarized intruders when data from days 4 and 5 were combined (although the difference was not significant when the two days were analyzed separately) (Suppl. Fig. 5A, C).

To determine whether the increased decoding accuracy was driven by the increase in aggression or the novelty of the intruder, we compared decoding accuracies when the same familiarized intruder was encountered on day 2, when aggression was high, and on day 3, when aggression decreased. When we tracked cells across days 2 and 3 and trained decoders to discriminate each behavioral dichotomy, we found no significant difference in accuracies across the two days (Suppl. Fig. 5F), suggesting that the enhanced decoding accuracy is not driven by differences in aggression alone in the absence of social novelty.

Next, we compared decoding accuracy on day 1, when the intruder was novel, but aggression was low (presumably due to the lack of aggressive experience by the resident), with decoding accuracy on day 2, when the intruder was familiar, but aggression was high (presumably due to the increased experience in attack of the intruder by the resident). There was no significant difference in decoding performance across these first two days (Suppl. Fig. 5G). This lack of change may reflect offsetting effects of the decrease in novelty and increase in aggression on day 2 compared to day 1.

### Increased behavioral decoding accuracy with novel conspecifics correlates with elevated social aggression but not identity processing through cross-day decoding

To examine whether the enhancement in dCA2 decoding of aggression towards a novel intruder was consistent with a causal role of dCA2 in promoting aggression to a novel intruder (see Fig. 2), we compared decoding accuracy of pairs of social behaviors with time spent engaged in a given behavior with familiarized and novel intruders in the different residents we examined. As the above decoding accuracy was obtained with equal, fixed lengths of behaviors across residents, the duration of each social behavior was normalized relative to the total time spent during the R-I test in minutes, thereby adjusting for combining behaviors on days 4 and 5. We found a significant positive correlation between the normalized duration of attack of a novel intruder with the accuracy of decoding attack behavior relative to exploration or dominance (Fig. 8C, right). The correlation for a novel intruder was specific for attack behavior as we found no significant correlation between either the normalized total duration of exploration or dominance behavior and attack decoding accuracy. In contrast to results with a novel intruder, we found no significant correlation between the normalized total duration of attack of a familiarized intruder and the accuracy of decoding attack behavior compared to either exploration or dominance behavior. The same trend for a positive correlation in attack and decoding accuracy was found without grouping days 4 and 5 (Suppl. Fig. 5E).

Finally, we investigated the stability of social behavior representations in dCA2 across two days of interaction with either the same familiarized intruder or the familiarized and novel intruder, using neurons tracked across sessions. Separate linear classifiers were trained to discriminate social behavior dichotomies on a familiarized day (either day 4 or 5 day based on maximal attack time). Decoding accuracy was then tested on a different familiarized day (day 3) or a novel day (day 6); the reverse training/testing protocol was then performed, and we computed the average decoding score (Fig. 8D). We found that cross-day decoding performed significantly better than chance on attack-related dichotomies across the two days of interaction with the familiarized intruder (Fig. 8E, left panel). In contrast, the classifier failed to decode above chance accuracy for behavioral dichotomies trained on the familiarized intruder and tested on the novel intruder, and vice versa (Fig. 8E, right panel). Since we trained on data from either day 4 or 5, depending on which day had greater duration of attack, and tested on data from days 3 or 6, in principle, a difference in cross-day decoding performance could reflect different intervals between the training and testing days. However, the maximal attack was most likely to occur on day 5 rather than day 4 (4/6 mice), so the interval between training and testing days was on average shorter with the novel and familiar mice (days 4/5 compared to day 6) than with the same familiar mouse (days 4/5 compared to day 3). These results are in agreement with our single cell correlation analysis (Fig. 6) and indicate that familiarized intruders are encoded through a stable representation while the neural code for social behaviors of a familiar and novel conspecific are more distinct.

## Discussion

In this study explored the relationship between the importance of dCA2 for SNR memory^2–4,18^ and its action to promote social aggression^19,22^. Our aim was to test the hypothesis that the mnemonic and aggression-promoting actions of dCA2 are closely related, positing that dCA2 provides a neural correlate of social novelty that specifically increased aggression towards a novel intruder. This hypothesis was based on three complementary sets of findings. First, prior studies^9,12,15,16,33^ and our present results (Fig. 1-3) show an increased probability of attack towards a novel relative to a familiar conspecific intruder. Second, dCA2 activity is required for social novelty recognition (SNR) memory, the ability of a mouse to discriminate a novel from familiar conspecific^2,4^. And third, both electrophysiological^24,25^ and cellular calcium imaging^26^ studies have demonstrated that dCA2 neural activity discriminates a novel and familiar conspecific, with a higher rate of spike firing around a novel compared to familiar conspecific^24,34^.

Using a longitudinally designed R-I test, we found that dCA2 does indeed promote aggression to novel intruders to a greater extent than to a familiarized intruder. Thus, chemogenetic silencing of dCA2 caused a greater suppression of attack towards a novel relative to a familiar intruder (Fig. 2B, D). Further support linking the role of dCA2 in SNR memory and aggression comes from the following finding shown in Figure 2B. Thus, when dCA2 was silenced during an initial encounter with a novel intruder not only was aggression suppressed but when that novel intruder was re-introduced the following day when dCA2 had recovered from silencing, the intruder provoked a high level of aggression as if it were being encountered for the first time. This suggests that when dCA2 was silenced, the resident failed to store a memory of the novel intruder. Surprisingly, silencing of dCA2 results in a lower level of aggression towards a novel compared to familiar intruder. This suggests that a distinct parallel circuit may be necessary for aggression towards a familiar conspecific.

Our calcium imaging experiments indicate that the role of dCA2 in promoting aggression may extend beyond social novelty recognition. Thus, dCA2 neuron activity, at both the single cell and population levels, discriminated the three classes of resident social behaviors towards the intruder of exploration, dominance and attack during the R-I test. Furthermore, population decoding of binary comparisons of exploration, dominance, and attack behaviors revealed significantly increased decoder accuracy towards novel relative to familiar intruder behaviors. Moreover, decoding accuracy was significantly correlated with time spent in attack of the novel but not the familiar intruder. Moreover, by examining changes in aggression towards a familiarized intruder, we found that the neural responses to the novel intruder are not simply a reflection of increased levels of aggression but rather are dependent on an interplay between intruder novelty and aggressive response.

Although Leroy et al. did not examine specifically an effect of intruder novelty, their fiberphotometry calcium recordings indicated an elevated activity in dCA2 in residents during bouts of exploration, dominance, and attack of intruders, with attack leading to the greatest increase from baseline. Our results, using longitudinally tracked single cell resolution of dCA2 pyramidal neurons, reveal that activity across the entire population of cells only moderately increases at behavior onset in a direct agonistic interaction. Surprisingly, we found that a greater proportion of dCA2 cells responded during exploration and dominance behaviors compared to attack. However, when we separated the entire population into activated and inhibited response types and aligned the responses to the onset of a given behavior, we found that neurons responded more vigorously to the onset of attack behaviors relative to exploration and dominance, with a greater mean transient calcium response, more similar to the results of Leroy et al.^22^. Our studies of the dynamics of neural activity revealed a further difference in how dCA2 responds to novel and familiar intruders as a novel intruder induced a greater time-locked increase in activity of attack-activated cells compared to a pre-attack baseline than did a familiar intruder. The apparent discrepancy between the results of Leroy et al.^22^ and our findings on mean population activity may be related to the recording method. Fiber photometry recording of calcium dynamics includes sub-threshold signals from the neuropil, in addition to the somatic suprathreshold responses^35^. Head-mounted miniscope calcium imaging allows offline segmentation of soma from background neuropil calcium activity facilitating the separation of sub-and suprathreshold responses and the longitudinal registration of cells across conditions^31,36,37^.

Additional findings by Leroy et al.^22^ demonstrate that dCA2 promotes social aggression through its excitatory outputs to dorsal lateral septum (dLS), activating dLS GABAergic interneurons that inhibit of a population of ventral LS (vLS) neurons that tonically inhibit VMHvl. In this manner dCA2 recruits a disinhibitory circuit that enhances VMHvl activity, thereby promoting aggression^22^. Accordingly, inactivation of dCA2 pyramidal neurons leads to a suppression of aggression by promoting tonic inhibition of VMHvl^22^. Importantly, this action of dCA2 appears to be under the control of the modulatory social neuropeptide arginine vasopressin. Thus, activation of the Avpr1b receptor, which is highly enriched in CA2 and has been implicated in both social memory and social aggression, by local AVP enhances the ability of dCA2 inputs to excite their dLS targets through presynaptic facilitation of glutamate release.

Two potential mechanisms may account for how dCA2 activity in response to a novel animal may promote aggression. First, enhanced firing observed in dCA2 pyramidal neurons in the presence of a novel animal^24,34^ may lead to increased output to the dLS, subsequently leading to greater disinhibition of VMHvl, thus triggering aggression. The second mechanism is based on results showing that a linear classifier can provide a generalized or abstract readout of novelty versus familiarity from the population level activity of dCA2 neurons^26^. Just as the classifier differentially weights neural activity to decode novelty from familiarity, so could a downstream neuron in dLS read out novelty from appropriately weighted dCA2 synaptic inputs^26^, firing in a biased manner towards novel conspecifics. This latter mechanism is supported by findings that the increase in dCA2 mean population activity towards a novel versus familiar conspecific can vary depending on experimental conditions^26^.

Several lines of evidence suggests that aggression is a malleable behavior capable of being learned and adapted to experience^9,11,38^. Indeed, the decision to engage in attack is likely influenced by several factors, including prior aggressive outcomes, mating experience, and the animal’s internal state at the time of an encounter. How might dCA2 integrate social novelty information to promote aggression?

Mice rely on olfactory information contained within the volatile fraction of urine from conspecifics for social identification and regulation of social behavior as chemical inactivation of the olfactory epithelium disrupts both SNR memory and aggressive displays^39,40^. The lateral entorhinal cortex (LEC) relays such social olfactory information from the olfactory bulb and olfactory cortex to dCA2 through both a direct projection and an indirect path through the dentate gyrus to CA3 to CA2 trisynaptic circuit. Moreover, dCA2 is capable of discriminating the volatile social odors contained in urine from male mice^27^ and silencing the direct LEC projection to dCA2 is sufficient to disrupt SNR memory^41^. Additionally, dCA2 receives modulatory projections from regions of the brain that relay social novelty cues, such as the supramammillary nucleus (SuM)^42^ and the medial septum (MS)^43,44^. dCA2 may integrate such information on social identity and social novelty with aggression signals provided by AVP inputs from hypothalamus. Whether AVP acts to selectively promote aggression towards a novel individual or acts indiscriminately to increase aggression remains unknown.

Several pathologies including Alzheimer’s disease (AD), autism, and other neurologic conditions are associated with both SNR memory deficits and heightened aggression^45^. Entorhinal cortex input into dCA2, necessary for the SNR memory function in rodents^41^, is especially high in neurofibrillary tangles^46^ in neurodegeneration progression, which are linked in AD aggression^47^.

Additionally, Avpr1b gene mutations have been linked to the aggressive behaviors seen in cases of childhood aggression, and autism^48,49^. Indeed, central AVP levels is correlated with history of aggressive behavior. In schizophrenia, patients showed a marked reduction in inhibitory but no change in excitatory pyramidal neurons^50^, further recapitulated in mouse models^51^. This loss of inhibition may lead to hyperactivation of dCA2, thus altering SNR processing and top-down regulation of aggression.

Together, our findings provide a mechanism by which SNR memory and social aggression interact as complementary processes within dCA2. Social novelty is integrated with social aggression signals in dCA2 and then translated into an adaptive response dependent on context, experience, and internal state. Our results suggest that dCA2 provides top-down regulation of aggression through its SNR memory function.

## Author contributions

A.V. and S.A.S. conceived the project and designed the experiments.

A.V. collected and analyzed the data with guidance from S.A.S. The data was interpreted by

A.V. and S.A.S. who wrote the paper.

## Data and Materials availability

All data and materials are available from the lead corresponding author with reasonable request.

## Acknowledgements

We thank L. Martinez and I. Nebo for assistance in obtaining immunofluorescent images and video annotations; D. Aranov, A. Bendesky, S. Hassan, L. Paninski, G. Petty, and W. Sheng for critical discussion and comments on the manuscript; T. Abe, L. Boyle, S. Fusi, and L. Posani for many valuable and technical discussion on analysis.

## Methods

### Key resources table

| REAGENT or RESOURCE | SOURCE | IDENTIFIER |
| --- | --- | --- |
| Antibodies |  |  |
| Rabbit polyclonal anti-PCP4 | Sigma-Aldrich | Cat #HPA005792; RRID: AB_1855086 |
| Mouse polyclonal anti-RFP | Invitrogen | Cat #710530; RRID: AB_2532732 |
| Bacterial and virus strains |  |  |
| AAV2/5.hsyn.GcaMP6f.WPRE.SV40 | Addgene (Douglas Kim & GENIE Project) | RRID: Addgene_100837 |
| AAV2/5-hSyn-hM4D(Gi)-mCherry | Addgene (Bryan Roth) | RRID: Addgene_50475 |
| AAV2/5-hsyn-mCherry | Signagen | Catalog #: SL116104 |
| AAV2/5.syn.jGCaMP8s.WPRE | GENIE Project | RRID: Addgene_162374 |
| Software and algorithms |  |  |
| Fiji | Schindelin et al. | https://fiji.sc/ |
| BORIS<sup>52</sup> | Friard and Gamba | https://www.boris.unito.it/ |
| CalmAn<sup>31</sup> | Giovannucci et al., 2019 | https://github.com/flatironinstitute/CalmAn |
| ANY-maze | Stoelting Co. | https://www.anymaze.co.uk |
| Python |  | https://www.python.org/ |
| Decodanda |  | https://github.com/lposani/decodanda |
| Inscopix Data Acquisition & Analysis Software | Inscopix | https://www.inscopix.com/ |
| Mesmerize | Kolar et al. 2021 | https://github.com/kushalkolar/MESmerize |

### Experimental model and subject details

All mouse procedures were performed in accordance with the NIH guidelines and with the approval of the Columbia University Institutional Animal Care and Use Committee. All mice used as residents in the resident intruder test were males on a C57Bl/6J background (The Jackson Laboratory). We used male BALB/cJ (The Jackson Laboratory) mice as intruders. Behavioral tests were performed on sexually naive male mice only. All mice were maintained on a 12-h light–dark cycle with ad libitum access to food and water. We used mice between 2 and 4 months old.

### Stereotaxic viral delivery

For imaging experiments, a volume of 200 nL AAV2/5.hsyn.GcaMP6f.WPRE.SV40 virus or AAV2/5.syn.jGCaMP8s.WPRE (respectively: titer: 1.3*10^12^ gc/mL; a gift from Douglas Kim & GENIE Project: Addgene viral prep # 100837-AAV5; http://n2t.net/addgene:100837 ; RRID:Addgene_100837; titer: 2*10^12^ gc/ml, gift from GENIE Project Addgene plasmid # 162374, virus packaged AAV2/5 in house; http://n2t.net/addgene:162374 ; RRID:Addgene_162374) was injected at a rate of 150 nL/min into the right hemisphere above dorsal hippocampal CA2 using stereotactic coordinates: (AP) - 2.0 mm, (ML) +1.8 mm, DV -1.2 mm (from dura) all from bregma. The pipette was retracted after 5 min.

In pharmacogenetic silencing experiments, we injected AAV2/5-hSyn-hM4D(Gi)-mCherry (titer: 1.05*10^12^ gc/ml; a gift from Bryan Roth (Addgene viral prep # 50475-AAV5 ; http://n2t.net/addgene:50475 ; RRID:Addgene_50475) or AAV2/5-hsyn-mCherry (titer: 2*10^12^ gc/ml) was bilaterally injected (200 nl per site) to express the inhibitory Designer Receptors Exclusively Activated by Designer Drugs (iDREADD) hM4Di or a fluorescent control in dCA2 using the same coordinates as above.

### GRIN lens implantation

Three weeks following viral injection, a 1.2 mm diameter circular craniotomy was centered at the following coordinates: AP -2.0 mm, ML +2.25 mm. We inserted a GRIN lens (Inscopix, 1.0 mm diameter, 4.0 mm length) into the craniotomy at a depth of -1.4 to -1.5 mm relative to bregma at a 10° angle from the midline, so that the lens was parallel to the CA2 cell body layer. The Inscopix Proview system imaged cells during implantation to adjust the position of the lens to optimize visible fluorescence. Kwik-sil was placed around the craniotomy and the lens secured in place using Metabond dental cement. The top of the Proview lens cuff was filled with lens paper and Kwik-cast to protect the lens. Mice were housed in pairs separated by a divider for one week before a plastic baseplate was placed over the lens and secured with Metabond dental cement. The baseplate and microscope were placed over the lens and the position was adjusted until cells were maximally in focus. After 1 week of recovery, mice were separated into their own home cages in preparation of acclimation to imaging setup.

### Immunofluorescent labeling and imaging

We perfused mice at the end of the experiments using saline followed by 4% PFA in ice-cold PBS. Brains were extracted and incubated in 4% PFA overnight. After 1 h washing in 0.3% glycine in PBS, 60-µm sections were prepared in coronal orientation using a Leica VT1000S vibratome. Sections were permeabilized and blocked for 1 hour with 5% goat serum and 0.4% Triton-X in PBS at room temperature. Sections were incubated overnight at 4°C in 0.1% Triton-X in PBS plus 5% goat serum. The following day, slices were washed three times for 10 minutes each in PBS and incubated with secondary antibodies for two hours. DAPI (ThermoFisher Scientific, #D1306) staining was applied at 1:4000 for 10 minutes in PBS at room temperature prior to mounting. Slices were mounted using Fluoromount (Sigma-Aldrich) and imaged using Zeiss LSM 700 confocal microscope.

For mCherry and PCP4 labeling, the first incubation was performed with mouse anti-RFP and rabbit anti-PCP4(respectively: 1:500, Invitrogen, # 710530 and 1:300; Sigma-Aldrich, #HPA005792, RRID: AB_1855086). The secondary incubation was performed with anti-mouse conjugated to Alexa 488 and anti-rabbit IgG2a conjugated to Alexa 568 (respectively: 1:500; #A21131, RRID: AB_2535771 and 1:500; #A11011, RRID: AB_143157).

For PCP4 labeling, the first incubation was performed with rabbit anti-PCP4 (1:300; rabbit anti-pcp4 #HPA005792, Sigma-Aldrich). The secondary incubation was performed with anti-rabbit conjugated to Alexa 568 (1:500; #A11011, RRID: AB_143157).

### Social aggression

The resident–intruder paradigm was used to assess social aggression as previously described^29^. Subject male mice (residents) were individually housed for a minimum of 1 week, with a cage change no less than 1 week before the encounter with a novel intruder.

Stimulus mice (male BALB/cJ intruders) were group housed and used for sequential encounters. Feeding and water apparatuses were removed for habituation to allow unimpeded interaction and better recording during testing. Ten-minute presentations (or 15 minutes during calcium recording) of weight-matched intruders occurred in the home cages of the resident mice after 30-minute habituation to the behavioral room.

Behavioral data was collected in a standard home cage arena measuring 18.5 cm x 38 cm, under red light illumination and sound attenuated conditions for later ethological analysis using Boris annotation software^52^. Videos were recorded perpendicular to the cage floor at 60 Hz using an Image source camera with a wide angle TPL 0420 6MP lens.

Ethological analysis of aggression was performed by a blinded observer. We measured: (1) the duration of attack; (2) the number of bites; (3) the duration of dominance displays; and (4) the duration of exploration displays. Operational definitions for these behaviors and their manual annotations are presented in Table 1.

**Table 1.** Operational definitions table.

| **Class** | **Description** | **Start Frame** | **Duration of Behavior** | **End Frame** |
| --- | --- | --- | --- | --- |
| **Attack** | Clear physical antagonistic interaction initiated by the resident mouse. | Resident is positioned and grounded for a bite characterized by outstretched forepaws and straight tail. Intruder displays submission in anticipation of an escape characterized by a sudden jolt post bite. | A bite can either be brief, marked by the intruder's immediate evasion or escape, or it can be longer, involving a sustained tethering action by the resident with a yanking motion. | Once the bite event is terminated. |
| **Exploration** | The resident mouse is engaged in (sometimes reciprocal) sniffing behavior towards the intruder, with the sniffing directed specifically at a body part of the intruder (usually anogenital region). | Once physical contact is made by the resident on intruder body. | Durations can be quite variable consisting of just a single sniff event or a prolonged contact with the intruder. | The departure of resident/intruder from the interaction. An exploration event can escalate to a dominance behavior. |
| **Dominance** | A clear imbalance in contact with the resident mouse manipulating intruder position. The intruder is usually still or anticipating an escape. | Resident is outstretched on intruder body by placing its forepaws on its hull or face. Mounting when the resident displays thrusting behavior. Tail rattle by resident. Chasing is often displayed by the resident when the intruder tries to evade a dominance episode. | Duration can be variable based on the type of dominance display. For example, an intruder tail rattle can be brief, while a mounting event can last several seconds. | The departure of resident from the interaction. The resident may deescalate to an exploration behavior or escalate to a full biting attack episode. |

### Pharmacogenetic silencing of dorsal CA2

Mice were injected with rAAV expressing iDREADD or inert fluorescent control and returned to their home cage for recovery. After 10 days of recovery the mice were housed singly. Habituation consisted of 5 days of Intraperitoneal (IP) Injection of saline prior to the social aggression test. On day 6, mice were injected with clozapine N-oxide (CNO) (5 mg/kg or 10 mg/kg in saline) or vehicle (saline) 30 min before testing. Each dose of CNO was analyzed separately and no significant difference was found using independent samples t-test between 5 and 10 mg/kg (t-stat = 0.83, p = 0.42) and so CNO data for both doses was combined.

A Linear Mixed Model (LMM) was employed to analyze the repeated measures R-I test data, with attack (or other social behavior) duration as the dependent variable. The model included fixed effects for Group (hM4Di: experimental or mCherry: control) and Day (1-6), their interaction, and a random intercept for each mouse to account for random variations between individual mice. The ‘mCherry:control’ group served as the reference category. The LMM was fitted using the Restricted Maximum Likelihood (REML) method. The analysis was conducted using the statsmodels package in Python. The model formula is as follows (using Attack as example):

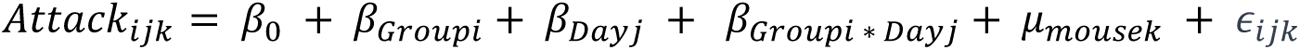

Here, *Attack_ijk_*, denotes the measured time spent in attack for the *i*th group on the *j*th day for the kth mouse, while β_0_ is the intercept of the model. β*_Groupi_*+ β*_Dayi_* represents the coefficients associated with different levels of the group variable or levels of day, respectively. β*_Groupi_*_∗_ *_Dayi_* denotes the interaction effects between the ‘Group’ and ‘Day’ variables. μ*_mousek_* accounts for intrinsic aggressive differences for each mouse *k*. ∈_*ijk*_ is the residual error term for each observation.

### Behavior with intruder experience

Mice were singly housed for 1 week prior to the start of the social aggression test. Subject (resident) mice were randomly assigned the condition of the corresponding intruder and aggression encounters were recorded for 10 minutes across 5 days. Three groups of residents were exposed to different types of intruders:

1. The ’familiarized’ group: These residents encountered the same intruder repeatedly throughout the experiment.
2. The ’naïve’ group: This group was introduced to a new intruder that they had never encountered before.
3. The ’experienced’ group: Residents in this group faced a unique intruder who was both new to them and had equal experience interacting as an intruder to other aggressive residents in an equal number of R-I tests.

A LMM was used assessed the fixed effects of ’Group’ (‘novel naïve’, ‘familiarized, and ‘novel experienced’) and ’Day’ (1-5), with each mouse accounted for as a random intercept to account for individual variability. The ‘familiarized’ group served as the reference category. The analysis included data from 8, 9, and 13 mice in Groups naïve, familiarized, and experienced, respectively. Significance of fixed effects and interactions was gauged using Wald tests with p-values reported.

### Data acquisition, preprocessing and motion-correction

On the day of the experiment, mice were moved to the behavior room. Mice were allowed to acclimate to the environment for 30 minutes. An nVista 3.0 Inscopix miniaturized microscope was inserted into the baseplate and used to record calcium fluorescence from dCA2 pyramidal neurons during social and non-social aggressive behaviors using Inscopix data acquisition software (20 frames per second, 50-ms exposure, 0.2-0.5 mW/mm2 EX-LED). The working distance between the microscope objective and the lens was adjusted to maximize cell focus, and this distance was maintained for all sessions. To align behavior and calcium videos, a 5V TTL pulse from an Ami-2 Optogenetic interface triggered calcium recordings through Anymaze software at the start of each trial along with a behavior video recording. Behavior recordings were collected at a rate of 60 Hz. The raw calcium videos were then run through Inscopix Data Analysis software for pre-processing as a concatenated habituation and test phase file. Videos were corrected for defective pixels and 4x spatially down-sampled. Background fluorescence was removed using a spatial band-pass filter and fluorescence videos were motion-corrected using the Inscopix motion correction algorithm. The preprocessed and motion corrected tiff files were then exported for cell identification and signal demixing (CNMFe) using the CaImAn package^31,32^.

### Segmentation and ROI identification

Cell regions-of-interest (ROIs) were identified using the Python CaImAn package for large-scale calcium imaging data. The spatial footprints and deconvolved signal for the active sources (ROIs) were extracted using CNMFe, and then the denoised, detrended temporal traces (over a 25 s window) and spatial footprints were exported. We used the Mesmerize package to evaluate individual ROIs and spatial footprints, and those with non-spherical or non-oval shapes caused by motion artifacts were excluded from analysis. Finally, the computed traces were normalized to obtain z-scores, dividing by the standard deviation of the raw fluorescence trace. The calcium data were aligned to the manually annotated behavioral movie based on the nearest time stamp, accounting for the time offset returned by anymaze relative to the onset of optical recording following an Ami-2 Optogenetic interface delivered pulse. We only included animals with stable window implants through all sessions.

### Longitudinal registration

For day-to-day longitudinal cell registration, we used a pipeline reported earlier^31^. Briefly, the approach takes a set of spatial components and finds components present in all using an intersection over union metric and the Hungarian algorithm for optimization. Cells were registered across days 3,4 and 5 and separately for days 4,5, and 6 to maximize cell retention on all days and form meaningful comparisons between familiarized-to-familiarized and familiarized-to-novel intruder representations.

### Behavior calcium recordings

Prior to the first test, mice were handled and habituated for six days on the following schedule: day 1: handling; day 2: handling, exposure to test room for 30 minutes; day 3: handling, exposure to test room for 30 minutes, and insertion of dummy miniscope overnight; day 4: handling, exposure to test room for 30 minutes; day 5: handling, exposure to test room for 30 minutes, exposure to test arena in home cage for 10 minutes, scruffing/insertion of the microscope, and removal of dummy miniscope; day 6: handling, exposure to test room for 30 minutes, exposure to test arena in home cage for 10 minutes scruffing/insertion of the microscope. In each test, subject mice were first allowed to habituate for 5 minutes in their home cage and interact with an intruder for 15 minutes. The same intruder mouse was placed in the subject mouse home cage over 5 days. On day 6 a novel and experienced intruder was placed in the same subject mouse home cage. Intruder mice were weight-and sex-matched to subject mice. In each session, the subject mouse was free to explore the arena. Boris software^52^ was used to manually score periods of interaction with the intruder consisting of non-overlapping exploration, dominance, or attack behaviors.

A LMM was used assessed the fixed effects of ’Day’ (1-6), with each mouse accounted for as a random intercept to account for individual variability. ‘Day 1’ served as the reference category. The analysis included data from 9 mice. Significance of fixed effects and interactions were gauged using Wald tests with p-values reported.

### Cell response

Neurons were classified as responsive for a particular behavioral variable (e.g. exploration, dominance, or attack) by comparing the activity during a given social behavior to activity during periods of non-social interactions during each R-I test. To assess statistical significance of each cell response, we computed a chance distribution from temporally shifted behavior labels^53^. Briefly, we employed a two-step procedure that maintained the neural calcium trace structure while destroying dependency with behavior labels, e.g. the animal’s behavior vector and neural traces. The temporal sequence of the behavioral vector was reversed by reindexing with flipped indices (e.g. the last index of the session became the first datapoint and vice versa). Subsequently, the flipped behavior vector was subjected to a random shift, where the shift magnitude, denoted as *n*, was randomly chosen within the range 0 ≤*n* ≤ the total length of the neural trace. For each shuffle the mean calcium rate was computed for time spent during a selected social behavior and non-social interaction, and the difference in mean calcium rates during these periods were recorded 1000 times, resulting in a null distribution. The true (non-shuffled) mean rate difference was compared to the shuffled distribution using a nonparametric two-sided permutation test. A neuron was deemed responsive for a behavior if the permutation test p-value was less than 0.05. If the rate difference was greater than 0, the cell was classified as ‘activated’; if the rate difference was less than 0, the neuron was classified as inhibited. A neuron that was not responsive for any behavior was considered ‘non-responsive’.

### Neural data correlation across behavior days

Neurons were tracked across the last 3 sequential days (4, 5, and 6) of the resident intruder test for direct comparisons (see section on longitudinal registration). The activity of each neuron (n = 311) was averaged over a 1-second interval, starting 500 milliseconds before and ending 500 milliseconds after the onset of each behavior. Spearman rank correlation (*r_s_*) was used to assess significance across each pair of behaviors and corrected for multiple comparisons using the Bonferroni method. Shuffling approach: For each behavior across days, cell IDs were independently randomized to disrupt any inherent association between the day variable, while preserving the overall structure of the dataset. Following the shuffling procedure, correlation coefficients were calculated for each pair of days within each behavior category, mirroring the analysis conducted on the unshuffled data. This step was repeated for 1000 iterations, creating a null distribution of correlation coefficients for each behavioral observation across days. The significance of the observed correlations was assessed by comparing them against the null distribution obtained from the shuffling process. A correlation value in the observed data was considered significantly different from chance if it fell outside the central 95% of the null distribution, corresponding to a p-value less than 0.05.

### Data labeling for decoding

For each subject (resident) and session (day), we selected neural data corresponding to periods in which the subject was actively interacting with the intruder in periods defined as non-social behavior, exploration, dominance, or attack (operational definitions in Table 1). This resulted in 3 social behavior dichotomies [exploration vs. dominance, exploration vs. attack, and dominance vs. attack] and 3 non-social behavior dichotomies. We then divided the conditions into binary dichotomies (class 0 and class 1) according to the variable we wished to decode.

### Cross-validation and pseudo-simultaneous population activity

Briefly, we divided data from each class of conditions (0 and 1) into training and test pseudo-trials, which each trial defined by a bout of interaction, with bout duration lasting from the beginning to end of a given interaction. Bout durations lasting longer than 500 ms were split into multiple 500-ms-long pseudo-trials. To avoid temporal artifacts, present in consecutive time bins through autocorrelation of calcium signals, contiguous trials were discarded prior to train/test splitting. We randomly selected 75% of pseudo-trials for training a classifier and the remaining 25% were used for testing decoding performance. The training data set was then used to train a SVM linear classifier, which was tested on the testing data set to assess the decoding performance. The whole procedure, from training-testing division to performance assessment, was repeated for *k*=10 times to implement a k-fold cross-validation scheme, taking the mean score (µ) as the estimated performance value of the decoding procedure. Decoding performances obtained by the cross-validated procedure were tested against a null model where the labels (0 and 1 as defined above) of pseudo-trials were randomly shuffled. After each shuffling, the same cross-validation procedure was repeated, obtaining a null-model value for decoding performance. We repeated the shuffling *n* times to obtain a distribution of null model performance values, yielding a mean null decoding performance (<µ>) and standard deviation of the null distribution (σ). The p value was then derived from the z-score of the performance computed on data compared to the distribution of *n* null-model values: z = [µ - <µ >]/σ.

### Cross-day decoding performance

Cross-day decoding for behavioral dichotomies was performed as previously described^26^. A separate linear SVM classifier was trained/tested on each behavior dichotomy (exploration versus dominance versus attack) for identity. Identity was chosen in pairs (familiarized-familiarized vs familiarized-novel). Cross validation and decoding performance were determined as defined in the ‘Cross-validation and pseudo-simultaneous population activity’ section.

## Figure Legends (Supplemental)

**Supplemental Figure 1:**
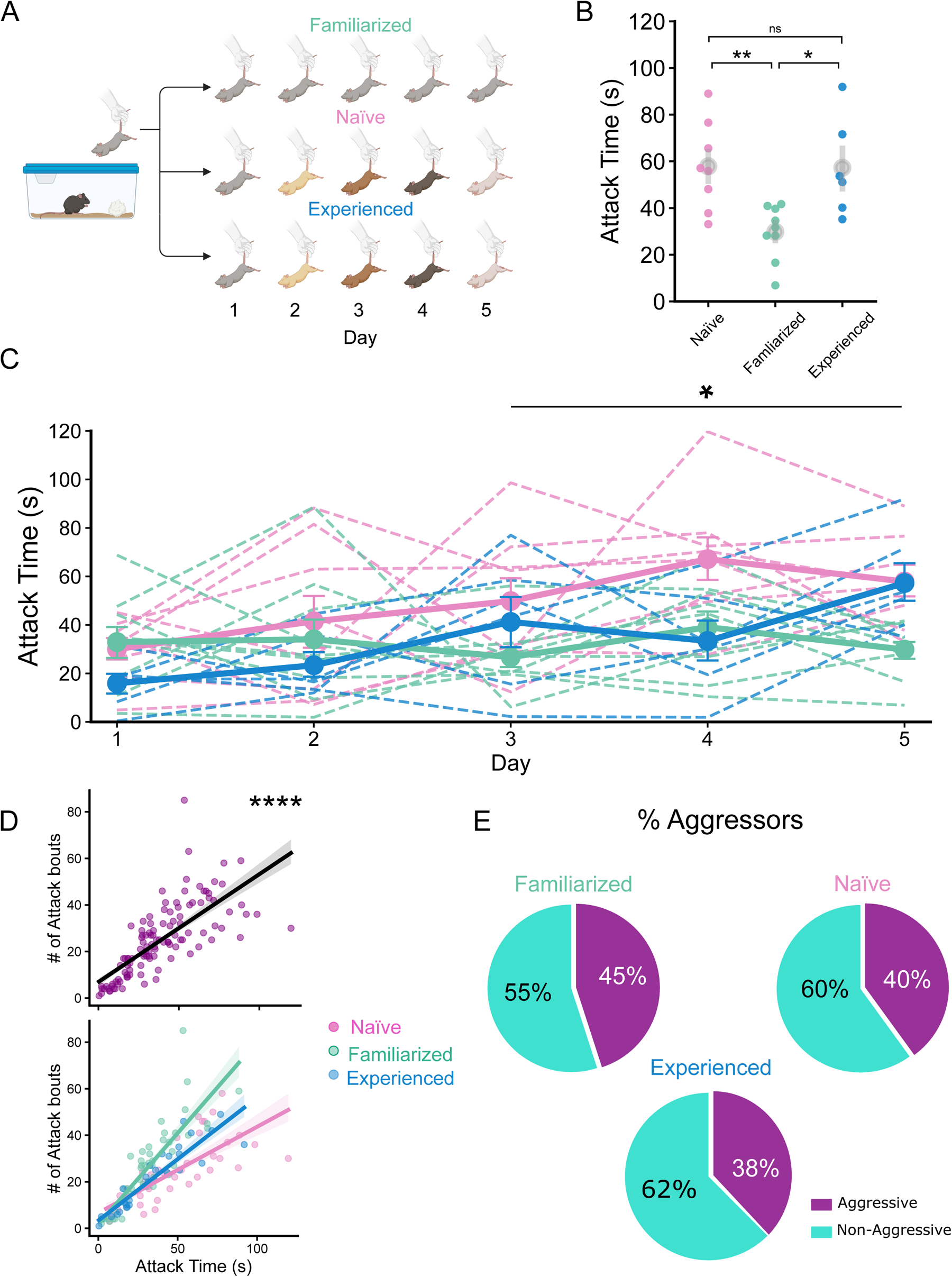
Social aggression increases to novelty even after accounting for intruder experience. **A)** Schematic and schedule of the 5-day modified resident intruder test for groups of resident mice with increasing familiar, novel- and naïve, or novel and experienced intruders across days. **B)** Resident mice in the familiarized intruder group show a significant decrease in time spent in attack on last day of test compared to residents in the novel naïve or novel experienced intruder groups. One way ANOVA, *F* (2, 20) = 7.19, *p* = 0.0013. Post hoc Tukey HSD test showed that the familiarized group alone differed significantly at p < 0.05 (0.0079, 0.017), from naive and experienced groups, respectively. **C)** Using a Linear Mixed Model (LMM), we analyzed attack durations across days for n = 9, 9, 6 mice in novel naïve, novel experienced, and familiarized (reference) groups. Interaction effects highlighted significant differences for the novel naïve group on Days 4 and 5 and for the novel experienced group on Days 3 and 5 (all p < 0.05) relative to the familiarized group. **D)** Time spent in attack (s) is correlated with total number of manually pressed attack bouts across groups. Top. For all groups, (Spearman’s rank correlation, *r_s_*= .83, *p* = 6.91e-31; n = 23 mice & 115 points). Bottom. For individual groups. Spearman’s rank correlation for familiarized group, *r_s_* = 0.88, *p* = 1.50e-15; n = 9 mice & 45 points, for novel naïve group, *r_s_* = 0.842, *p* = 9.79e-12; n = 8 mice & 40 points, for experienced novel group*, r_s_* = 0.95, *p* = 2.57e-15; n = 6 mice & 30 points. No significant difference between correlations: Fisher z-transformed values of familiarized vs. novel naïve: -0.668, *p* = 0.503, novel naïve: vs. novel experienced: -2.26, *p* = 0.024, and familiarized vs. novel experienced: -1.71, *p* = 0.088. **E)** Proportion of mice that were aggressive or not in each group as measured by the presence of at least 1 biting episode across all groups and used in the analysis. No significant difference in proportions between groups (*χ*2 statistic: 0.22, *p* = 0.90).

**Supplemental Figure 2:**
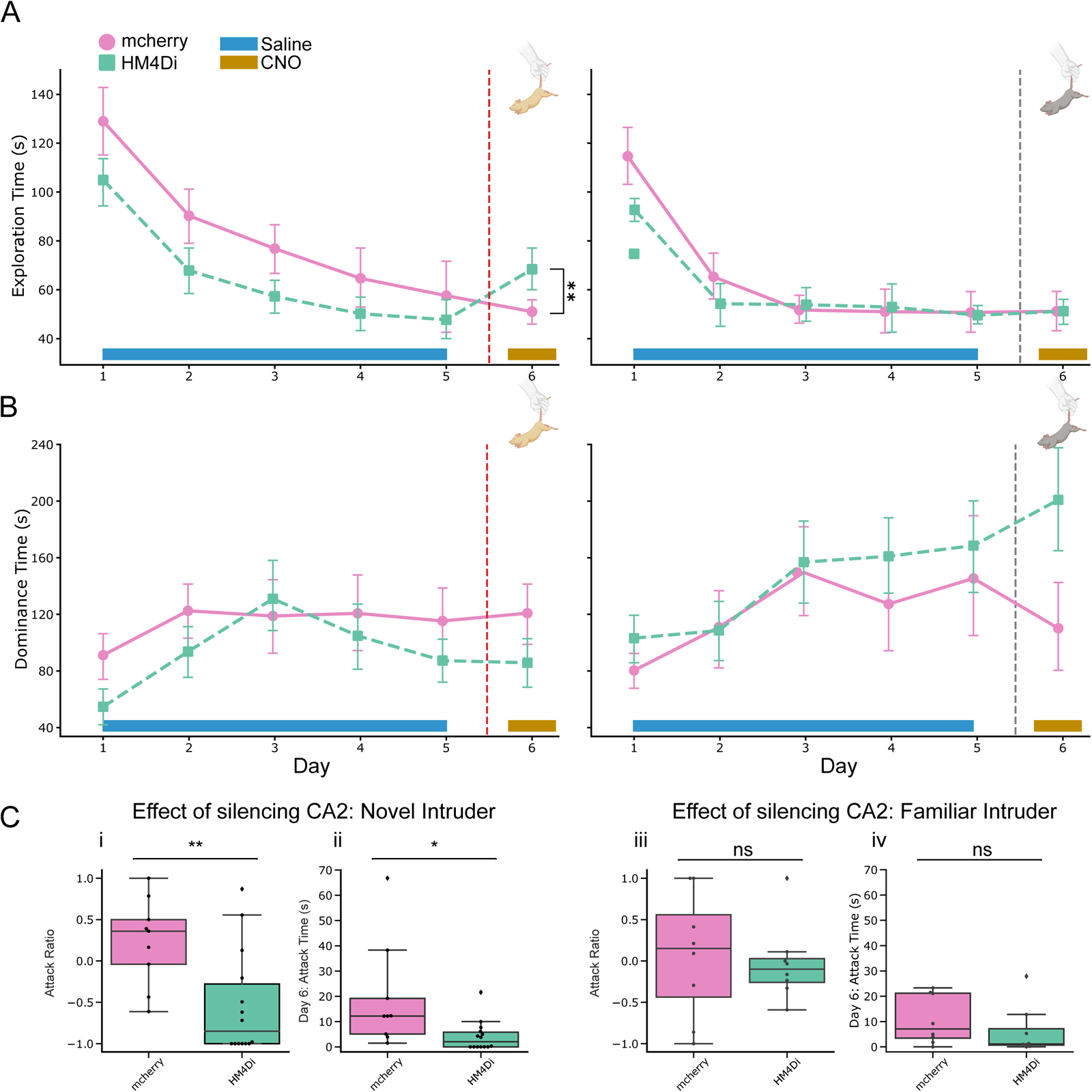
Silencing of dCA2 across social behaviors and days. **A)** Time spent in exploration for mice in control and experimental groups across 6 days. Experimental: iDREADD, control: mCherry (Left Panel-Novel Day 6) Experimental (n = 14) and control mice (n = 9) show a significant difference in exploration time on day 6 (LMM; interaction effect for day 6, p = 0.009). (Right Panel-Familiarized Day 6) Experimental and control mice (n = 8) show no difference in the time spent in exploration behaviors across days. **B)** Time spent in dominance for mice in control and experimental groups across 6 days. (Left Panel-Novel Day 6) Experimental (n = 14) and control mice (n = 9) show no difference in the time spent in dominance behaviors across days. (Right Panel-Familiarized Day 6) Experimental and control mice (n = 8) show no difference in the time spent in dominance behaviors across days. **C)** Attack Ratio: (Day 6 – Day5)/ (Day6 + Day5) and absolute attack time on day 6. i. Attack ratio shows a significant difference between control-novel and experimental-novel (paired sample t-test: *t-stat* = 2.52, *p* = 0.0067), and ii. significant difference in absolute attack time (paired sample t-test: *t-stat* = 3.01, *p* = 0.0199). iii. No significant difference between experimental-familiarized and control-familiarized on attack ratio (paired sample t-test: *t-stat* = -0.937, *p* = 0.754) or iv. absolute attack time (paired sample t-test: *t-stat* = -0.318, *p* = 0.364).

**Supplemental Figure 3:**
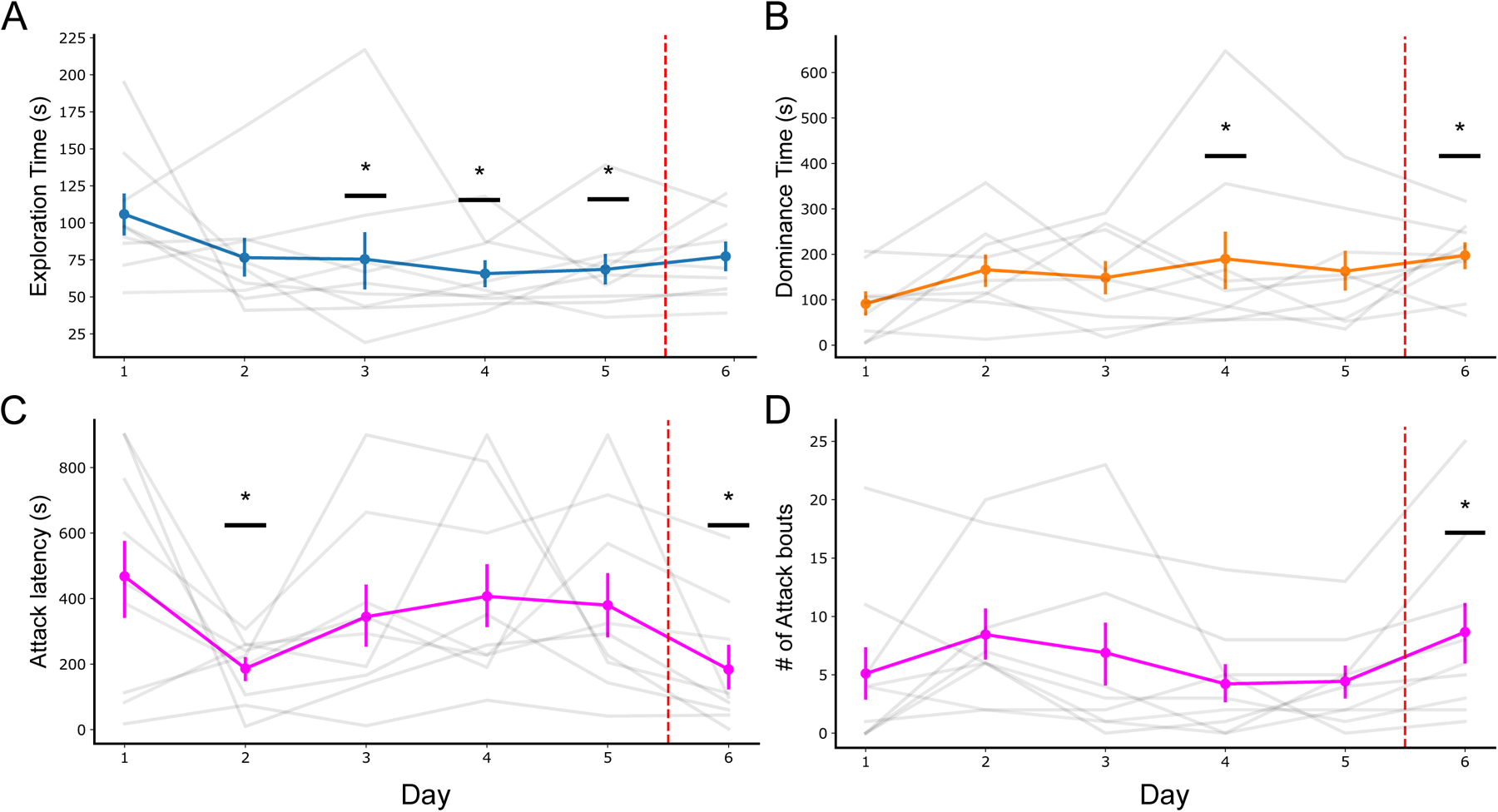
Social behaviors of imaged resident mice. **A)** Time spent in exploration for all imaged aggressive mice across days (n = 9). Day 3,4 and 5 showed a significant different (LMM-with day 1 as reference, p-values (p <.05), Day 3: 0.05, Day4: 0.01, Day5: 0.016. **B)** Time spent in dominance for all imaged aggressive mice across days (n = 9). Day 4 and 6 showed a significant different (LMM-with day 1 as reference, p-values (p <.05), Day4, *p* = 0.028; *p* = Day 6: 0.018). **C)** Latency to first attack for all imaged aggressive mice across days (n = 9). Days 2 and 6 demonstrated significant decrease in attack latency (LMM-with day 1 as reference), Day2, *p* = 0.013; Day6, *p* = 0.012. **D)** Number of attack bouts for all imaged mice across days (n = 9). Day 6 alone showed a significant difference (LMM-with day 1 as reference, p-values (p <.05), Day6: .047).

**Supplemental Figure 4:**
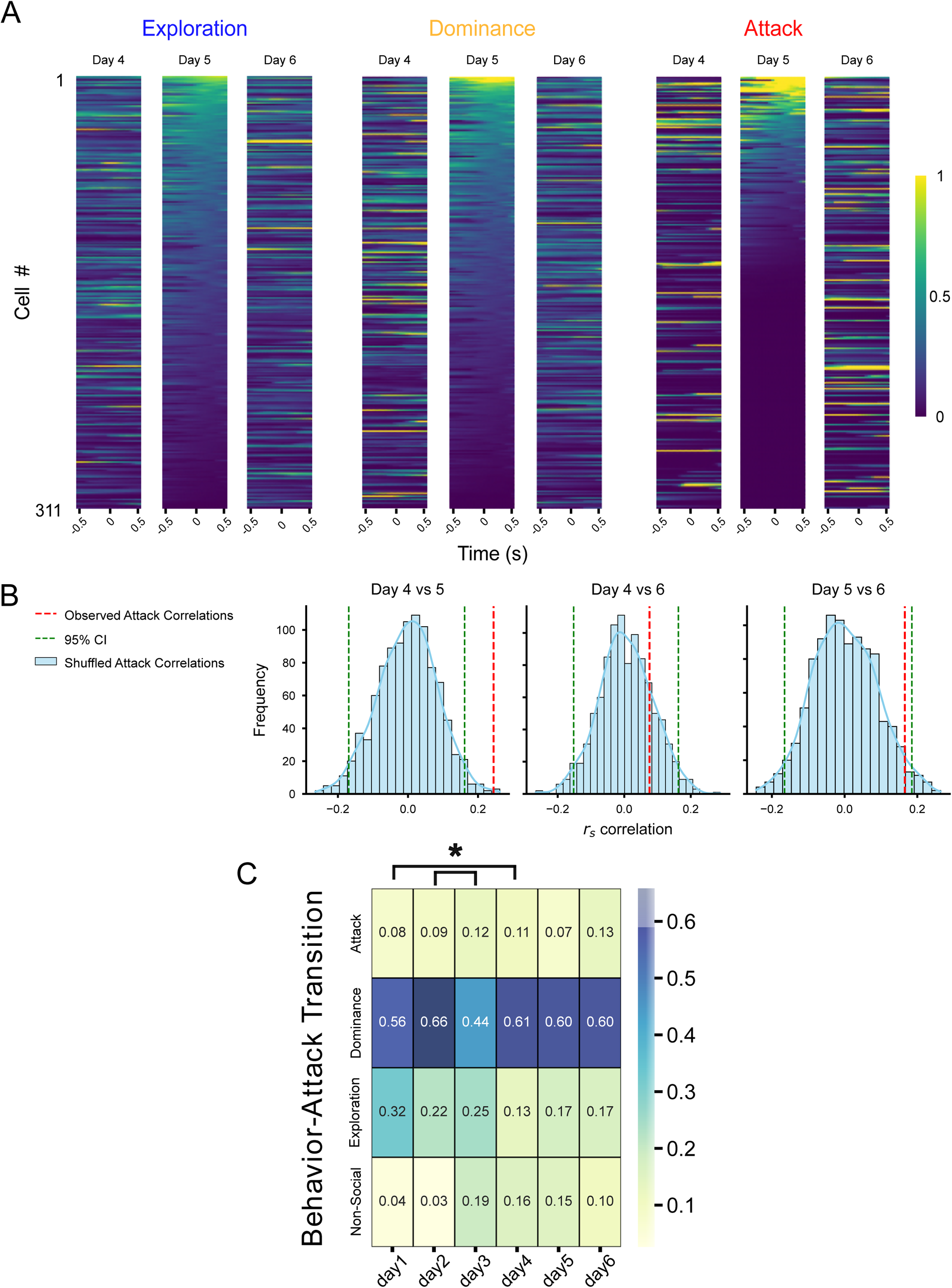
Correlation of neural activity and behavior transitions. **A)** Average neural vectors for 311 from n = 6 aggressive mice on days 4, 5, and 6 aligned to behavior onset bounded by 500 ms windows. Neurons are corresponding across rows and sorted based on day 5 mean activity from 0 to 500 ms after behavior onset. **B)** Shuffling cell IDs for each behavior across days independently. For each pair of days, a null distribution of correlation to test significance of observed value (red dashed line). Day 4 vs 5, *p =* 0.006; Day 4 vs 6, *p* = 0.676; Day 5 vs 6, *p* = 0.088. **C)** Behavior transition matrix reporting transition of behavior to attack. A behavior taking place 3 seconds prior to an attack bout is reported, otherwise a ‘non-social’ behavior class is reported.

**Supplemental Figure 5:**
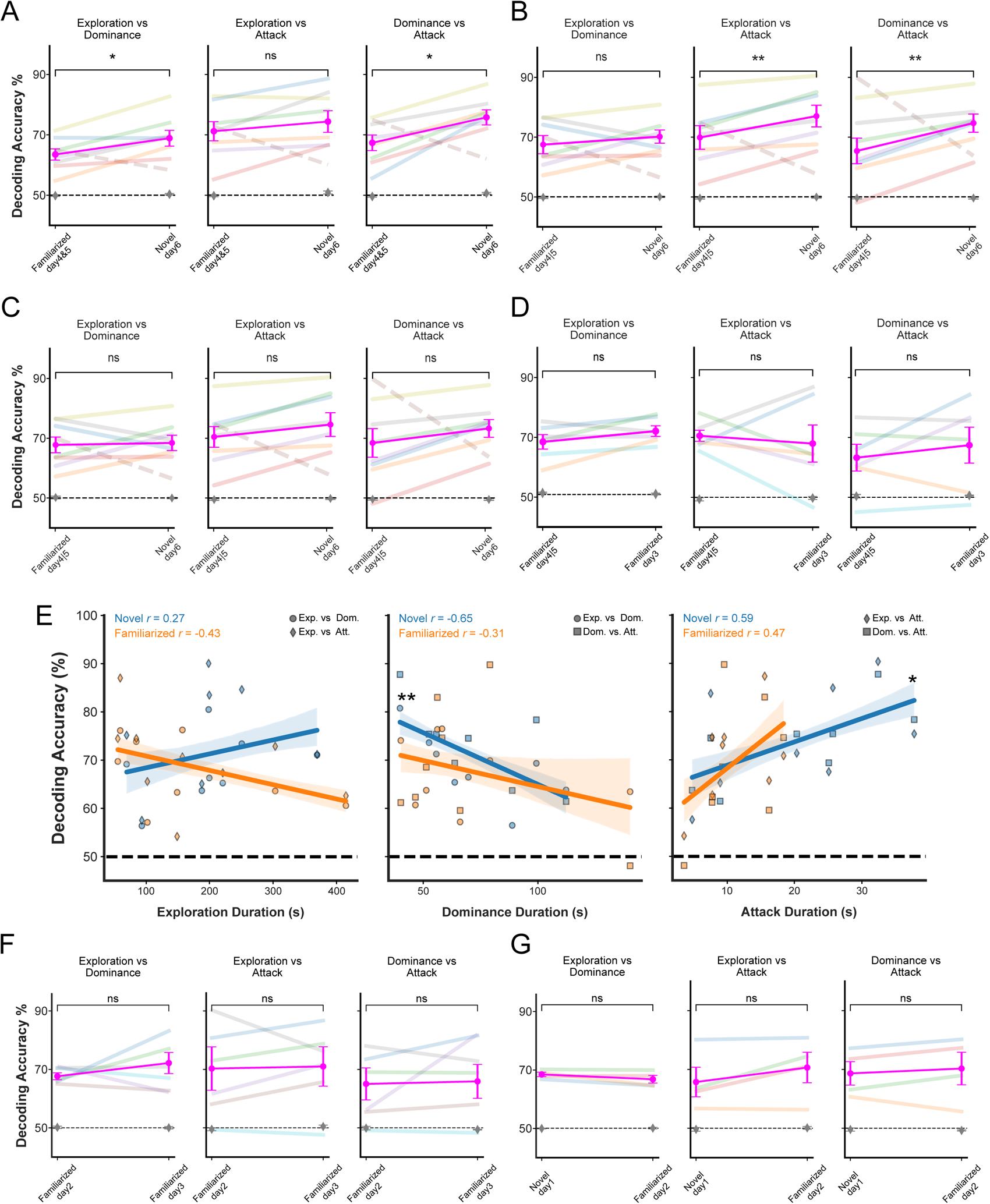
Decoding on familiar days and controlling for experience. **A)** Decoding accuracy of behaviors with the novel intruder on day 6 compared to the interactions with the familiarized intruder using combined data from days 4 and 5 with inclusion of aberrant mouse. Exploration versus dominance, *t-stat* = 2.41; *p* = 0.046, exploration versus attack, *t-stat* =1.13; *p* = 0.29, and dominance versus attack, *t-stat* = 2.42; *p* = 0.046 (paired t-tests; n= 8). **B)** Decoding accuracy of behaviors with the novel intruder on day 6 compared to the interactions with the familiarized intruder using data from maximal attack time on day 4 or 5. Exploration versus dominance, *t-stat =* 0.916*; p* = 0.395; exploration versus attack, *t-stat =* 4.74; *p* = 0.00316; and dominance versus attack, *t-stat =* 5.87*, p* = 0.00107 paired t-tests; n = 7; data from one aberrant mouse that showed decreased attack to the novel versus familiarized intruder was excluded from analysis). **C)** Decoding accuracy of behaviors with the novel intruder on day 6 compared to the interactions with the familiarized intruder using data from maximal attack time on day 4 or 5 with inclusion of aberrant mouse. Exploration versus dominance, *t-stat* = 0.202; *p* = 0.845, exploration versus attack, *t-stat* = 1.25; *p* = 0.252, and dominance versus attack, *t-stat* = 1.05; *p* = 0.328 (paired t-tests; n= 8). **D)** Decoding accuracy of behaviors with the familiarized intruder on day 3 compared to the interactions with the familiarized intruder using data from maximal attack time on day 4 or 5. Exploration versus dominance, *t-stat* = 2.22; *p* = 0.0771, exploration versus attack, *t-stat* = -2.328; *p* = 0.0674, and dominance versus attack, *t-stat =* 0.789; *p* = 0.465 (paired t-tests; n = 6). **E)** Decoding accuracy for indicated behavior dichotomies plotted against duration (seconds, s) a resident was engaged in exploration (left), dominance (middle) or attack (right) of familiarized (orange) or novel (blue) intruders. Pearson’s correlation coefficient is reported. Exploration versus dominance or attack: Novel intruder, *r* = 0.27, *p =* 0.311; familiarized intruder, *r* = -0.43; *p =* 0.092); Dominance versus attack or exploration: Novel intruder, *r* = -0.65, *p =* 0.006); familiarized intruder, *r* = -0.31, *p* = .23); Attack versus exploration or dominance: Novel intruder, *r* = 0.59, *p =* 0.015; familiarized intruder, *r* = 0.47, *p =* 0.063. **F)** Decoding accuracy of behaviors with the intruder on day 2 compared to the interactions with the familiarized intruder on day 3. Exploration versus dominance, *t-stat* = 0.574; *p* = 0.591, exploration versus attack, *t-stat* = 0.622; *p* = 0.561, and dominance versus attack, *t-stat* = 1.11; *p* = 0.316 (paired t-tests; n = 6). **G)** Decoding accuracy of behaviors with the intruder on day 1 compared to the interactions with the intruder on day 2. Exploration vs. dominance, *t-stat* = -1.622; *p* = 0.203, exploration vs. attack, *t-stat* = 1.748; *p* = 0.179, and dominance vs. attack, *t-stat* = 0.732; *p* = 0.517 (paired t-tests; n = 4).

